# CHIMERA-DDR: A Machine Learning Framework for Classifying Heterogeneous Mismatch-Repair and Homologous-Recombination Deficiency Patterns in Prostate Cancer

**DOI:** 10.1101/2025.10.08.680958

**Authors:** Komal Sharma, Divin A. Wilson, Yu Wang, Shabi Haider, Vineel Bhatlapenumarthi, Daniel Boiarsky, John M. Cipriaso, Leo Yermakov, Rajanya Dutta, Manavita Vashisth, Ilsa M. Coleman, Armand Bankhead, Shantanu K. Jadhav, Bhabishya Neupane, Bradley W. Taylor, Colm Morrissey, Michael. T. Schweitzer, Robert B. Montgomery, Sridhar Rao, Marja T. Nevalainen, Ariel A. Nelson, Emmanuel S. Antonarakis, Patrick G. Pilié, Jacob E. Berchuk, Kevin K. Zarrabi, Colin C. Pritchard, Anai Kthari, Hui-Zi Chen, Ben George, Razelle Kurzrock, Gavin Ha, Peter S. Nelson, Paul L. Auer, Anjishnu Banerjee, Deepak Kilari, Navonil De Sarkar

## Abstract

Current DNA damage repair (DDR) biomarkers employ binary classifications that fail to capture the molecular complexity of tumors with concurrent repair deficiencies. We used genomics analysis to stratify 672 metastatic prostate cancer patients into 11 DDR subgroups, identifying 51 molecular signatures with weighted roles in class identity. We identified a tumor-mutational-burden very-high subset, characterized by 19 mutations/Mb or more, as a molecularly distinct group characterized by preserved genomic integrity and enhanced immunogenicity. Critically, 2.3 percent of tumors exhibited concurrent TMB-High and HRR mutant phenotypes, while 1.5 percent harbored MMR bi-allelic loss without MMRd (mismatch-repair-deficiency) signatures. Clinical validation in 130 patients demonstrated superior immunotherapy responses in tumors with very high TMB levels. We developed CHIMERA DDR, a probabilistic machine learning tool that integrates these 51 genomic features using a nested Random Forest architecture to infer seven clinically relevant DDR subgroups. After negating model overfit concerns, CHIMERA-DDR showed exceptional classification performance (AUCs 0.919-0.999) to accurately detect MMRd and HRR mutant molecular subtypes with or without concurrent DDR deficiencies, resolving admixed phenotypes to enable precision therapeutic stratification beyond binary methods.

## INTRODUCTION

Prostate cancer (PC) is projected to cause over 313,000 new diagnoses and nearly 36,000 deaths in 2025 in the United States (1). While early-stage disease is curable, metastatic prostate cancer (mPC) remains incurable with a poor prognosis. Precision medicine has improved outcomes through targeted therapies, particularly PARP inhibitors (PARPi) for homologous recombination repair-deficient (HRRd) tumors and immune checkpoint inhibitors (ICIs) for mismatch repair-deficient (MMRd) cancers (2–7).

PARPi targeting homologous recombination repair-deficient (HRRd) tumors, particularly those with BRCA1/2 mutations, have demonstrated notable efficacy, and treatment guidelines now recommend somatic molecular testing to stratify mPC patients for PARPi therapy. Approximately 45% of BRCA2-mutant PC patients fail PARPi therapy despite harboring the canonical predictive mutation (8–10). Similarly, ICI responses vary widely in biomarker-selected populations. Pembrolizumab is approved for tumors with high tumor mutational burden (TMB ≥10 mutations/Mb), microsatellite instability-high (MSI-H), or for mismatch repair deficient (MMRd) molecular subset (7,11–15). Biomarker-selected patients demonstrate improved but variable therapeutic outcomes, yet only 16% of TMB-high cancers exhibit MMR deficiency or high MSI (16–25) While TMB ≥20 mut/Mb correlates with better response rates (∼58% vs. ∼20%), this trend isn’t uniform across cancers (14,23–31).

Despite these multimodal advances, current clinical decision-making still relies on a binary interpretation of DNA repair activity, assuming either a singular or a homogeneously admixed concurrent repair-deficient phenotype (e.g., concurrent HRR gene mutation and MMRd with TMB-High state). However, even dual biomarker-positive tumors frequently fail to respond to their corresponding therapies, underscoring the complexity of DNA repair biology and the limitations of current biomarker strategies(32–34). Binary approaches cannot resolve intratumor heterogeneity or the functional interrelationships between concurrent genomic events - a major gap limiting therapeutic stratification. Current paradigms fail to capture DDR pathways’ complex interrelatedness, dynamic nature, and phenotypic variability driven by clonal heterogeneity and variable genomic aberration penetrance (e.g., BRCA1 C61G hypomorphic variants) across evolving phenotypic landscapes (35–37).

Consequently, there is a critical need for dynamic, multimodal tissue biomarkers that address tumors’ molecular and functional heterogeneity by deconvoluting functional deficiency spectra within clonally admixed phenotypes of varying degrees, capturing how concurrent DNA repair gene mutations with variable penetrance shape overall tumor phenotype and guide precision treatment strategies in advanced PCs (Supplementary Fig. 1, Graphical abstract). To address this challenge, we performed deep curation of DNA repair pathway genes. We analyzed genes from various repair pathways, including DNA break sensors (ATM and CHEK2, among others), DNA mismatch repair pathway genes, Homologous recombination repair pathway genes, and other DDR regulatory genes such as ATR, AR, and RecQ helicases (38). The genomic features were analyzed together with a comprehensive battery of functional signature scores derived from DNA and RNA sequencing data. This resulting integrated approach refined the molecular classification of DNA repair subgroups within established clinical stratification frameworks for advanced PC.

We delineated molecular subgroups along the DDR axis in a discovery mPC cohort for more precise classification and validated this refined substructure in an independent advanced PC case series. Finally, we utilized this comprehensive understanding to develop a probabilistic modeling framework that accurately classifies advanced PC patients based on their functional DNA repair activity profiles (Supplementary Fig. 1). This model deconvolves DNA repair status into components, representing divergent admixed phenotypes based on their relative importance in overall phenotype, while estimating both the degree of phenotypic heterogeneity and the dominant phenotype within PC tumor samples. The platform leverages multiple orthogonal functional readouts to infer the relative contributions and interactions of co-existing DNA repair deficient phenotypes (e.g., HRRd, MMRd) as well as DNA repair intact compartments. Such multidimensional approaches may be particularly valuable in developing precision biomarkers for context-aware therapeutic stratification.

## RESULTS

### Fine-Scale Molecular Stratification of mPC via TMB Thresholds and DDR Alterations

We first assessed the prevalence of tumors with high tumor mutational burden (TMB-H; ≥10 mutations/Mb) in advanced PCs. Whole-exome sequencing of 672 tumors (228 from University of Wisconsin Tissue Acquisition Necropsy(UW-TAN) case series, 444 from Stand Up to Cancer(SU2C) case series) yielded 482 high-quality exomes after rigorous filtering (Supplementary Table 1). Of these, 44 tumors (∼9.1%) met the TMB-High criterion (≥10 mutations per Mb), an FDA-approved biomarker for selecting pembrolizumab treatment in solid tumors (7). Notably, 33 of the 44 TMB-High tumors lacked detectable mutations in HRR, RecQ, and DSB sensor pathway genes, although a subset of these tumors harbored MMR pathway deficiencies (n=14, ∼42.4%). Such an enriched case set provided a robust foundation for characterizing this molecular subtype.

To delineate the substructure of TMB-High tumors, we subclassified the TMB-High tumors into DDR proficient and DDR-deficient mPC. This analysis encompassed 46 curated DDR-related genes across Homologous recombination repair (HRR), Mismatch repair (MMR), Fanconi anemia, Double strand break sensor (DSB Sens.), RecQ helicase pathways, and a few core DDR regulators (Fig. 1A-B). We examined tumor ploidy, percent genome alteration (PGA), genomic scar scores (LOH, LST, TAI, percent LOH), microsatellite instability (MSI), and mutational signature profiles. These analyses revealed patterns within TMB-High tumors and further subclassification across DDR pathways (Fig. 2A-F, Supplementary Fig. 2A, Supplementary Table 2A-D, Supplementary Table 3A-B). Importantly, as expected, tumors with relatively higher TMB were prominently enriched for MSI-High score and COSMIC MMRd signature positivity (Fig. 1B). To identify the optimal TMB cut-point maximizing MSI-High score enrichment, we applied binary logistic regression with three-fold cross-validation (Supplementary Fig. 2B-E), identifying 18.61 mut/Mb as the most discriminative threshold. The goal of this analysis is to enrich MMRd tumor samples to better characterize this molecular subgroup using multimodal molecular approaches. Tumors exceeding this cutoff were designated as TMB-very high (TMB-vH), distinct from the redefined TMB-High (TMB-H) class encompassing the 10-18.61 mutations/Mb range (Supplementary Fig. 2 E). Based on this specific cutoff, approximately 3.9% of this case series qualified as TMB-vH. Overall, 1.2% of the entire case series were characterized as TMB-H with concurrent HRR pathway loss-of-function mutations (TMB-H HRRm), and 1.03% were TMB-vH with HRR loss-of-function mutation (TMB-vH HRRm) where we defined HRRm class tumors with detected HRR pathway gene mutations (Molecular group definitions are in Supplementary Document 1). Molecular profiling of this relatively rare subset revealed a molecular characteristic dichotomy that splits the samples into two distinct subgroups: i) MMR-deficient-like tumors (MMRd; characterized by very high TMB, MSI-High status, MMRd signature positivity, and low ploidy/PGA/HRD-Scar scores), ii) HRR deficient-like tumors (HRRd; exhibiting lower TMB/MSI but elevated ploidy, PGA, and HRD-Scar scores), independent of MMR pathway gene loss status, likely reflecting distinct tumor evolutionary trajectories (Fig. 2A-F, Supplementary Table 2A-D, Supplementary Table 3A-B, also refer to Supplementary Document 1 for HRRm & HRRd definition).

**Fig. 1:**
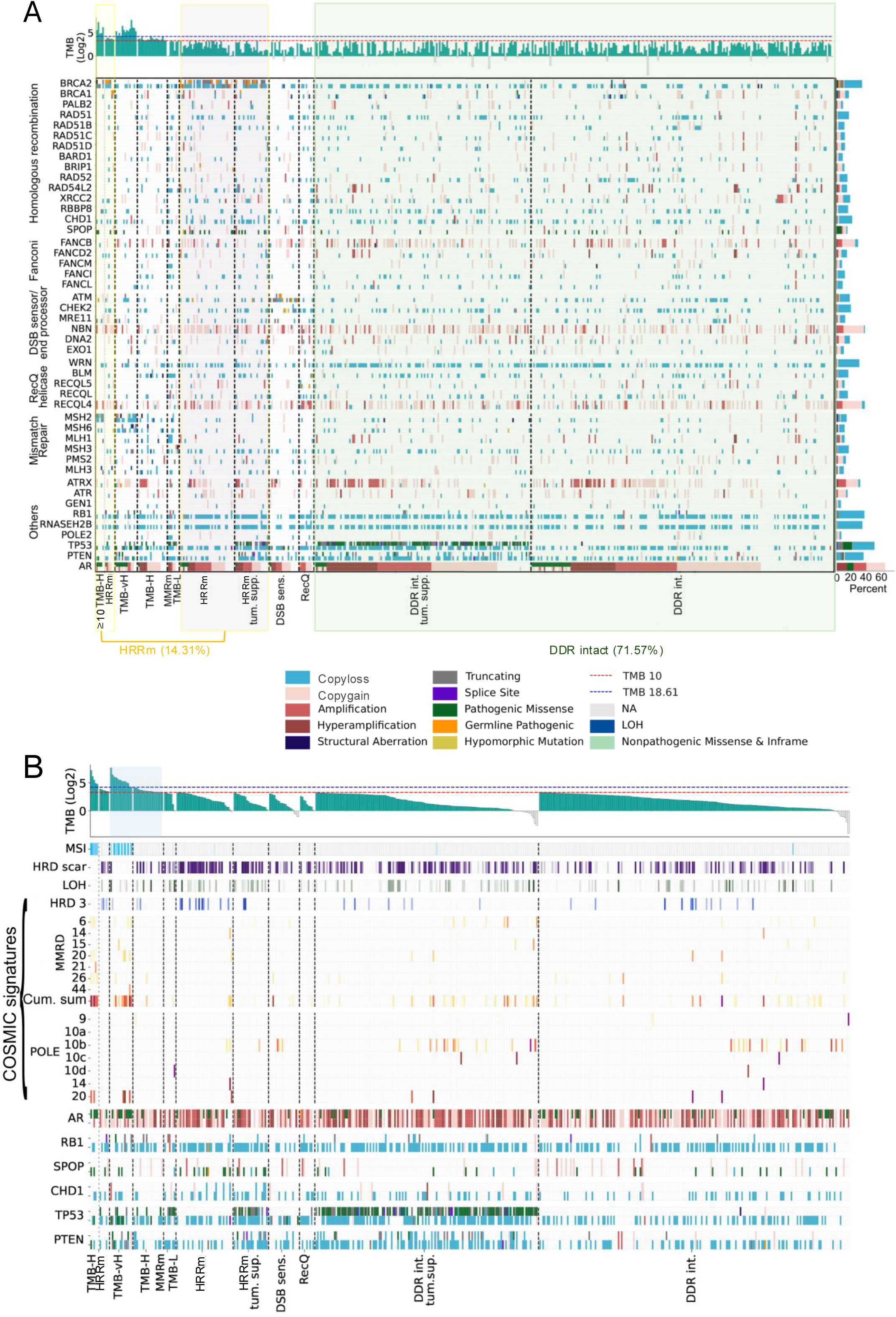
Genomic Stratification of Advanced Prostate Cancer Reveals Distinct Molecular Substructures Across DDR Pathways. (**A**) Curated genomic calls and tumor mutation burden analysis revealed 14.31% of advanced prostate cancers are HRR bi-allelic mutant, 2.3% harbor both TMB-High (>10 mut/Mb) and HRR mutations, while 71% are DDR-intact. We identified ∼3.9% of advanced prostate cancers as TMB-vH using our 18.61 TMB cutoff, with this subset enriched for MMR pathway gene bi-allelic loss-of-function mutations. The HRR subset showed the highest BRCA2 loss-of-function frequency. (**B**) Using the 18.61 TMB cutoff revealed substantial molecular dichotomy within the TMB-H subgroup, distinguishing TMB-vH tumors with concurrent HRR mutations. All TMB-vH tumors, regardless of concurrent HRR mutation status, were enriched for MSI-High scores, AR pathogenic mutations, and COSMIC MMRd signatures while negatively enriched for AR copy amplification and HRD-Scar signatures(p=0.76). The TMB-High 10-18.61 subgroup was instead enriched for HRD-Scar scores. Based on observed signatures, we detected no distinctive molecular differences between TMB-vH and TMB-vH with concurrent HRR mutations. Copy number annotations were performed using the homebrew tool pyTHiA-CNA (Supplementary Document 5).

**Fig. 2:**
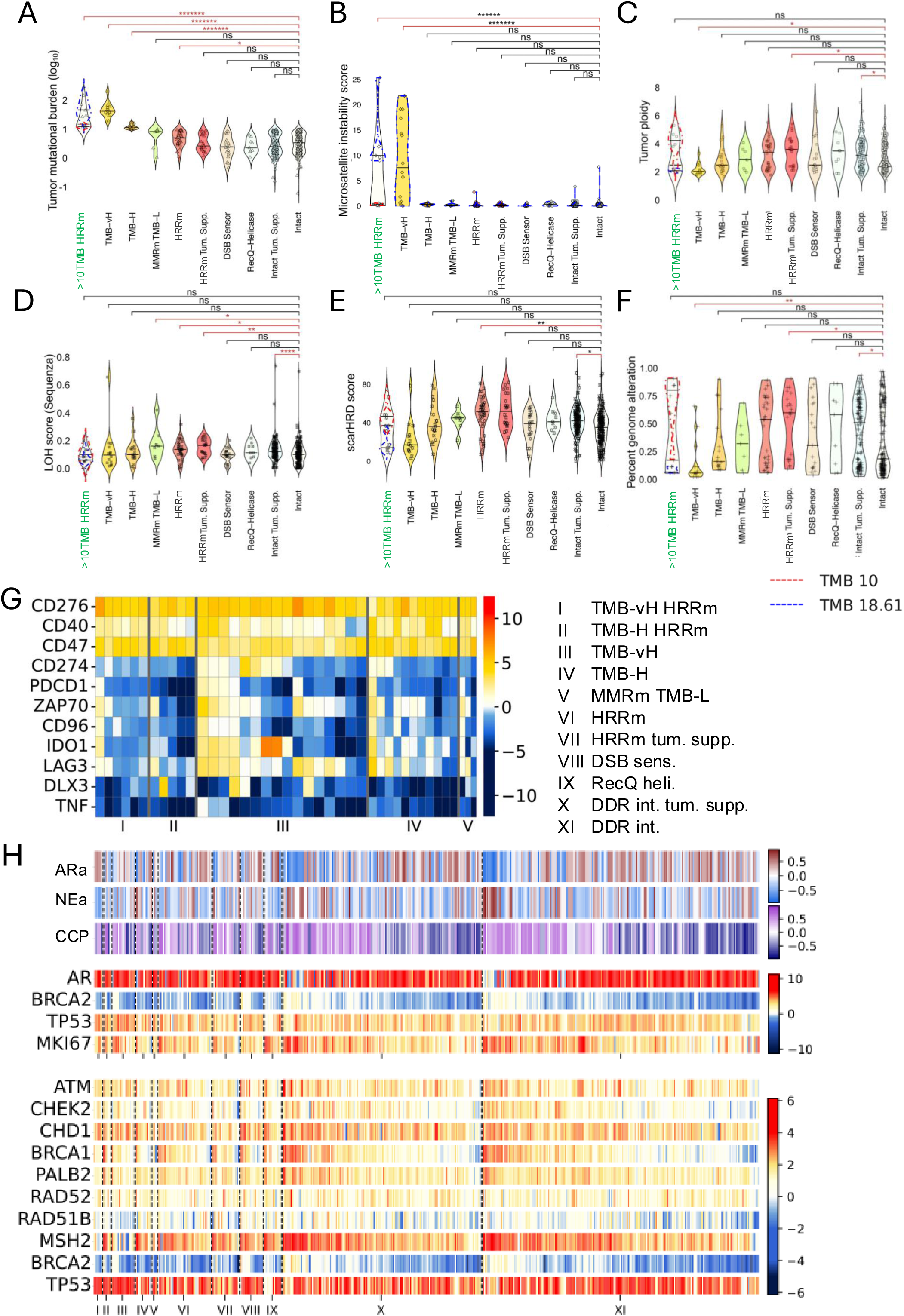
Multi-Dimensional Molecular Profiling Enables Fine-Scale Classification into 11 Molecular Subclasses Along the DDR Axis Through Integrated Genomic, Transcriptomic, and Pathway-Specific Signature Analysis. (**A-F**). Distinctive intrinsic molecular feature dichotomy within concurrent TMB-High and HRR mutant subsets clearly supports the molecular basis for 18.61 TMB cutoff-based subgrouping (Wilcoxon rank test). Analysis revealed molecular dichotomy based on tumor mutation burden, microsatellite instability score, tumor ploidy, HRD-Scar score, and percent genome alterations. LOH scores from Sequenza showed no striking differences. We report significant differences in tumor ploidy, LOH, HRD-Scar score, and PGA between DDR-intact and DDR-intact subgroups with bi-allelic tumor suppressor gene mutations (Wilcoxon rank-sum test for continuous variables like TMB, ploidy, PGA and LOH) (Fisher’s exact test for categorical features like MSI and Scar, *::p<0.01, **:: p<0.001, ***::p<0.0001, ****::<0.00001, *****::<0.000001, ******::<0.0000001, *******::<0.00000001). All other comparisons shown as ‘ns’ stand for statistically non-significant. HRD-Scar). Note, 4 (TMB-vH HRRm, TMB-H HRRm, TMB-vH & TMB-H) of the 10 subgroups are defined by TMB status. Significant P-value is expectedly high or can be ignored. PGA analysis was performed using home brew tool PGA_Calculator [https://github.com/Niel20/PGA_Estimator, Supplementary Document 6] (**G**) Gene expression profiling revealed ∼50% of TMB-vH subclass tumors exhibit elevated PD-L1 and LAG3 expression. (**H**) TMB-vH subgroups demonstrated consistently elevated TP53 mRNA expression (corrected p-value = 0.005). TMB-vH HRR mutant subgroups displayed elevated AR activity scores with low NE scores but high CCP scores, contrasting with TMB-H subgroups that showed reduced AR activity despite elevated CCP scores. BRCA2 expression exhibited strong correlation with MKI67 and CCP scores across most subgroups, though this relationship was disrupted in TMB-vH-HRR mutant tumors (Pearson’s correlation analysis, ρ= −0.351). MSH2 and RAD51B showed notably low expression enrichment in TMB-vH subsets regardless of HRR mutation status, maintaining consistently above-median expression levels in TMB-vH tumors while other subgroups displayed broader dynamic expression ranges.

Building on prior findings (39,40), which refer to bi-allelic TP53 or PTEN alterations to elevated HRD-Scar and LOH scores even in HRR-wild-type tumors, we further refined the classification of HRRm and DDR-intact tumors by TP53 and PTEN mutant statuses to better align biological mechanisms with clinical relevance. Ultimately, these analyses precisely defined 11 molecularly distinct DDR axis subgroups within the mPC case series, forming the basis of a novel stratification approach that informs future therapeutic evaluation and tool development (Fig. 1A-B).

### Preserved Genomic Integrity and Heightened Immunogenicity Define TMB-vH Tumors

Comparison of TMB-vH (≥18.61 mut/Mb) and TMB-H (10 - 18.61 mut/Mb) tumors revealed extensive molecular divergence (Fig. 2A-F). TMB-vH tumors were enriched for bi-allelic MMR gene mutations, especially in MSH2 (Fig. 1A; Supplementary Fig. 2F-G). 10 out of 14 tumors exhibited the COSMIC MMRd signatures (Fig. 1B). Such enrichment was notably absent in TMB-H tumors. Despite their high mutational burden, TMB-vH tumors exhibited lower ploidy (p = 0.008) and reduced PGA (p = 0.01) and lacked highly recurrent arm-level alterations (e.g., 8p loss, 8q gain, 17p loss), which are typical of other mPC subtypes, including HRR gene mutant subsets (Fig. 2C, 2F; Supplementary Fig. 3A-B). These findings indicate that genomic structural integrity is preserved despite hypermutation.

Immune infiltrate frequency analyses (Total n=323, SU2C n=236; UW-TAN n=87) using CIBERSORT (v1.02) revealed that the TMB-vH HRRm subset had the highest mean and median CD8⁺ T cell infiltration, while the TMB-H HRRm group had overall reduced immune cell density (Supplementary Fig. 5A). TMB-vH tumors exhibited a trend toward increased CD274 (PD-L1) mRNA expression (p=0.006, q=0.06) and LAG3 downregulation (p=0.02, q=0.27), though neither reached statistical significance after multiple testing correction. (Fig. 2G-H, Supplementary Fig. 4B-C, Supplementary Table 4A-B) (n=467); further supporting enhanced checkpoint inhibitor sensitivity. Recent clinical case series-based outcome aligns with these findings: patients with TMB > 20 mut/Mb and MMR deficiency have shown improved responses to ICI (14).

### Enrichment of AR Pathogenic Mutations and Disrupted Molecular Dependencies in TMB-vH Tumors

To further evaluate alterations in mPC driver events with respect to their frequency in tumors with the relatively intact genomes found in TMB-vH tumors, we investigated androgen receptor (AR) alterations across mutational burden subsets. The AR is commonly altered either by mutation or locus amplification in mPCs following AR pathway targeting. TMB-vH tumors were significantly enriched for pathogenic AR ligand-binding-domain (LBD) mutations associated with ARPI resistance yet notably lacked AR focal amplifications - a pattern consistent with the genome structural integrity defining this subgroup (Fig. 1A; Supplementary Fig. 5A-B). Among all molecular classes, TMB-vH exhibited the highest frequency of AR pathogenic mutations driving selective resistance to ARPI inhibitors (Supplementary Fig. 5B). Survival analysis & PSA90 objective response analysis showed no significant differences in outcomes between AR mutant and AR wildtype tumors within the TMB-vH subgroup, indicating that conventional AR mutation based prognostic models may not be applicable to this molecularly distinct subset (Supplementary Fig. 5C, Supplementary Fig. 6A-B).

To determine whether TMB-vH tumors exhibit altered gene expression and growth regulatory dependencies, we evaluated established proliferative and aggressive phenotype markers across all 11 DDR axis subgroups, including cell-cycle progression (CCP) score, neuroendocrine activity (NEa) score, MKI67 and BRCA2 mRNA expressions, and androgen receptor activity (ARa) score (41–43)(Fig. 2H. Across the majority of molecular subgroups, we validated established prognostic associations, wherein AR activity exhibited strong inverse correlations with NE activity, CCP score, MKI67 expression, and BRCA1/2 mRNA expression levels (Supplementary Table 5A-C). These patterns were consistent with prior evidence linking elevated CCP scores to NE-driven aggressive biology and inverse relationships with AR activity (Supplementary Table 5D-F). (42,43). In contrast, the TMB-vH HRRm & TMB-vH subgroup displayed a complete inversion of these canonical relationships in terms of mRNA expression as well (Fig. 2H, Supplementary Table 5A-B). Elevated CCP scores were decoupled from NE activity and instead co-occurred with high AR activity, elevated BRCA2 expression, and increased MKI67, representing fundamental disruption of established prognostic marker interdependencies (Supplementary Table 5E-F). This molecular signature analysis indicates that TMB-vH HRRm tumors operate under a distinct regulatory program that diverges entirely from the NE-associated proliferative phenotype defining aggressive behavior in other mPCs.

Comparative analyses of TMB-H versus TMB-vH tumors identified significantly reduced frequency of 8q gain in TMB-vH tumors, with this chromosomal arm alteration being consistent with their compromised genomic structural integrity (Supplementary Fig. 3A-B, Supplementary Table 6A-B). However, within the TMB-vH HRRm subset, we identified a profound disruption of these canonical molecular relationships, which fundamentally challenges the current understanding of proliferative regulation in hypermutated tumors (Supplementary Fig. 4A-B).

We also identified a previously unrecognized molecular subtype defined by biallelic mutations in MMR pathway genes that paradoxically exhibits low TMB (< 10 mut/Mb, MMRm TMB-Low) (44). This subgroup demonstrated a distinctive genomic characteristic, with a very high frequency of 8p loss and 8q gain involving RAD21, along with MYC amplification. Notably, this subgroup also harbors evidence of focal MYC amplification and frequent AR hyper-amplification (Fig. 3G, Supplementary Fig. 3A-B, Supplementary Fig. 5A). Unlike TMB-vH tumors, the median tumor ploidy is approximately 3 (p=0.03), with a very high average LOH score (Fig. 2C-D, Supplementary Table 6C), greater than even the HRRm subset of the mPC group median LOH (9.5) (Supplementary Table 6D) (15). Surprisingly, despite these genomic alterations, this subgroup does not show significant statistical deviation in HRD-Scar signature scores from HRRm or HRRm-tumor suppressor groups (MMRm TMB-L 45 vs. HRRm 51.5 (p=0.18) or HRRm tum. supp. 52 (p=0.31), numbers represent group median HRD-Scar score) (Supplementary Table 6E). The key distinguishing feature that differentiates this subgroup from HRRd tumors is the proportion of genome altered (PGA, MMRm TMB-L 0.31 vs. HRRm 0.53 (p=0.27) or HRRm tum. supp. 0.59 (p=0.13)), highlighting the complex interplay between different DNA damage repair pathways and their molecular characteristic manifestations (p-values were non-significant due to high PGA score variance across HRRm samples) (Fig. 2A-F, Supplementary Fig. 2A-D, Supplementary Table 6F).

**Fig. 3:**
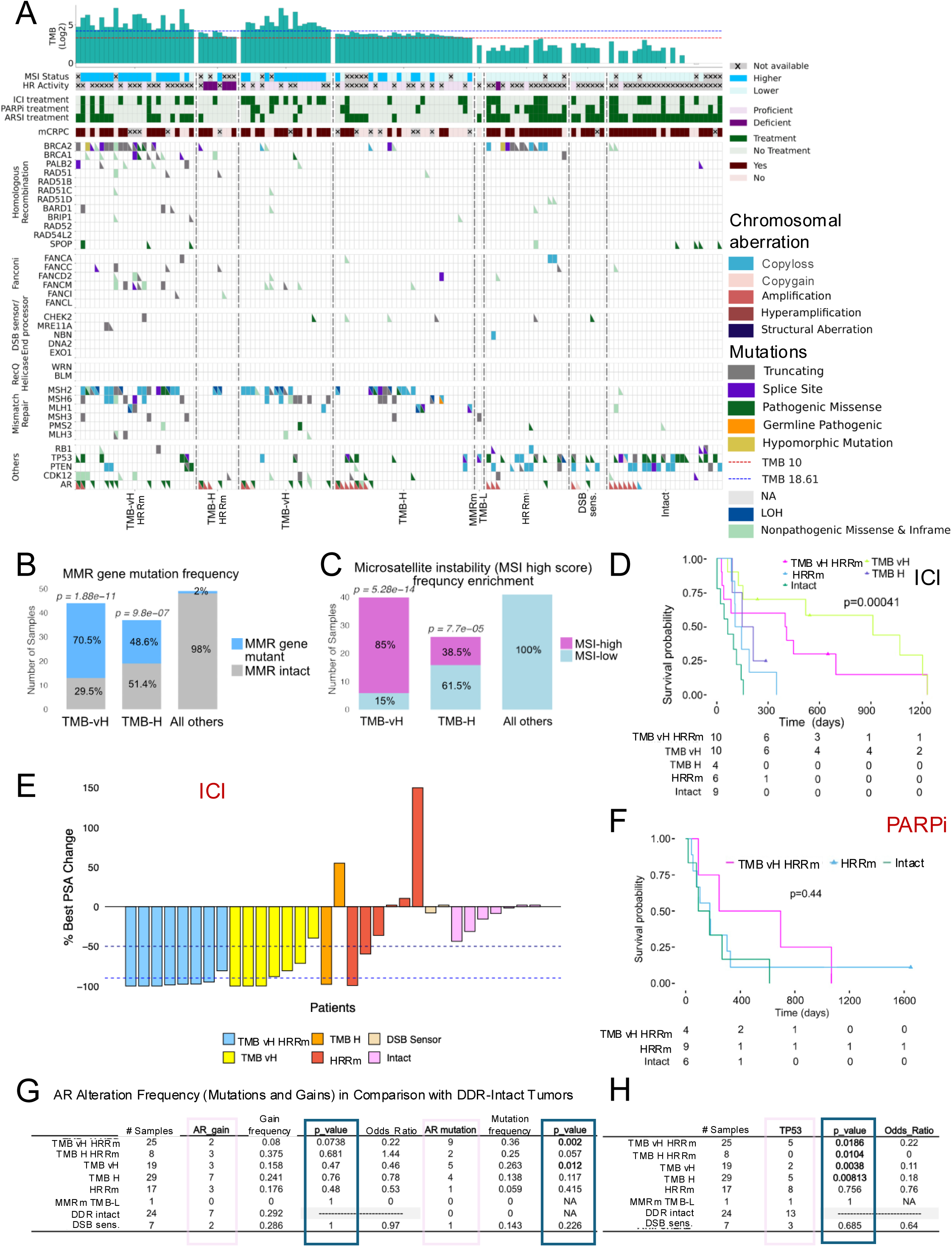
Clinical Cohort Analysis Validates TMB >18.61 Cutoff Defining a Distinct MMRd-Like Molecular Subset with Enhanced ICI Response in Advanced Prostate Cancer. (**A**) Heatmap demonstrates substantial molecular subtype representation in real-world pooled case series from 4 institutions. This cohort shows overrepresentation of concurrent TMB-H or TMB-vH tumors with or without HRR mutations due to targeted patient selection for robust validation, and substantial underrepresentation of DDR-intact subsets. Since MSI scores and detected mutations are clinical-grade assays that validate our discovery cohort-derived molecular phenotype patterns, this bolsters accuracy of our proposed molecular subgrouping. Heatmap is plotted with home brew tool D-AV plotter (Supplementary Document 7) (**B**) Enrichment of MMR pathway gene bi-allelic mutations in TMB-vH subsets independent of concurrent HRR mutation status validates discovery findings. (**C**) MSI score enrichment in TMB-vH subsets confirms inference accuracy observed in the primary 482-sample discovery cohort. (**D**) Survival curves demonstrate TMB-vH and TMB-vH HRR mutant subsets show significantly superior immune checkpoint inhibitor response, highlighting clinical relevance of accurate TMB-vH subtype determination in prostate cancer. (**E**) Swimmers plot reveals 100% of TMB-vH HRR mutant and 85% of TMB-vH tumors achieved PSA50 response—significant enrichment compared to HRR mutant or DDR-intact subsets (p<0.005, Fisher’s exact test). (**G**) Consistent with discovery cohort findings, we observed significant AR pathogenic mutation enrichment, though unlike the discovery cohort, negative enrichment of AR copy amplification events was not statistically significant. (**H**) Mutation frequency enrichment analysis revealed borderline enrichment of detected TP53 mutation events in all four TMB >10 mut/Mb groups with and without concurrent HRR mutations compared to DDR-intact subgroups.

### Intragroup Heterogeneity in Molecular and Functional Signatures Leads to Two Distinct Molecular Classes in Concurrent TMB-High and HRR Mutant Tumors

Within the mPC case series, we identified a rare subset (2.3%) harboring both high tumor mutational burden (TMB >10 mut/Mb) and bona-fide loss-of-function mutations in homologous recombination repair genes (TMB-H HRRm), a molecular type previously described in isolated case reports and one single study using a clinical genome (32–34,38). This dual-alteration group exhibited two clearly separable molecular profiles that paralleled the broader TMB-H population dichotomy. The TMB-vH HRRm subclass (≥18.61 mut/Mb) displayed features consistent with MMRd, including consistent MSI-High status, enrichment for COSMIC MMRd mutational signature scores, and comparatively lower tumor ploidy, percent genome alteration (PGA), and HRD-Scar scores (Fig. 1B, Fig. 2C-E). This MMRd-like subclass was enriched for AR ligand-binding domain mutations, mirroring the TMB-vH profile observed in MMRd tumors, and exhibited high CD8⁺ T-cell infiltration, suggesting an immunologically active phenotype (Supplementary Fig. 4A, Supplementary Fig. 5A).

Critically, the TMB-vH HRRm subset demonstrated the same inversion of canonical prognostic marker relationships observed in TMB-vH tumors: the AR activity score showed a reversed correlation trend with CCP score (ρ =0.53, p=0.28), complementary with the established negative correlations observed across other molecular subgroups (Fig. 2H, Supplementary Fig. 4C, Supplementary Table 5C). In contrast, the TMB-H HRRm subclass (10–18.61 mut/Mb) resembled a functional HRRd phenotype, characterized by lower MSI-High frequency but markedly elevated HRD-Scar scores, PGA, and tumor ploidy, providing functional evidence of HRRd-like molecular signature feature (Fig. 2A-F).

Tumors with genomic alterations reflecting both HRRd and MMRd consequently feature biomarkers predicting high response rates to two distinct targeted therapeutic strategies: ICI (driven by high TMB and MMRd-like features in TMB-vH HRRm) and PARPi (driven by functional HRRd in TMB-H HRRm). However, the pronounced variation in genomic and phenotypic features within this dual-alteration subset suggests therapeutic responsiveness may differ substantially depending on the dominant underlying biology, underscoring the critical need for precise molecular stratification to optimize treatment selection.

### Clinical Validation Case Series Recapitulates Molecular Features and Reveals Therapeutic Significance of DDR Subtype Stratification

To assess the clinical relevance of the mPC DDR genomic subtypes identified in initial cohort of metastatic tumors, we assembled an independent validation case series comprising genomic data derived from 130 men with mPC and attendant clinical outcomes. Based on available clinical sequencing interpretations and molecular signatures, we defined eight clinical subgroups within a targeted validation case series of 130 patients’ data points (Supplementary Table 7). This case series intentionally overrepresented with TMB-vH HRRm (19%) and TMB-vH (19%) tumors, enabling robust assessment of these rare molecular subtypes using TMB thresholds derived from our discovery analysis and clinical-grade sequencing reports from established diagnostic providers (Fig. 3A-H). The clinical case series recapitulated key molecular features of TMB-vH tumors: significant enrichment for MSH2 mutations and AR ligand binding domain pathogenic mutations, with notable lack of genome wide copy number gains and amplifications consistent with preserved genomic integrity (Fig. 3G-H, Supplementary Fig. 6C). Importantly, we observed significant negative enrichment of TP53/PTEN loss-of-function events in the TMB-vH subgroup (Supplementary Fig. 6C). Unlike the discovery case series, the clinical TMB-H subgroup contained substantial numbers of patients with bi-allelic loss of MMR pathway genes (p=9.8e^−07^) or MSI-High status(p=7.7e^−05^), though at frequencies lower than the TMB-vH subset (Fig. 3B-C).

Assessments of the genomic subclassification with clinical outcomes in this cohort revealed divergent therapeutic outcomes that underscore the clinical significance of TMB-vH molecular stratification. While TMB-vH tumors in the discovery case series trended toward poorer prognosis with limited ICI or PARPi exposure, both TMB-vH and TMB-vH HRRm subgroups demonstrated improved overall survival in the validation case series, likely reflecting higher frequency ICI and PARPi utilization (Supplementary Fig. 6D-E). TMB-vH and TMB-vH HRRm tumors showed significantly superior ICI response rates (Kaplan-Meier analysis and PSA50 response) (Fig. 3D-E, Supplementary Fig. 6F). Notably, TMB-vH HRRm patients exhibited prolonged ICI+PARPi combination therapy (Fig. 3F), whereas no differential response to androgen receptor signaling inhibitors (ARSI) was observed across subgroups (clinical case series p=0.74) (45,46) (Supplementary Fig. 7A-B).

### Dynamic Prediction of the Relative Composition of DDR Pathway Defects Using CHIMERA-DDR: A Probabilistic Model for Precision DDR Stratification

The genomic analyses of tumors with DDR alterations indicated that current binary DDR classification approaches fail to capture the full molecular complexity of these tumors, particularly MMR deficiency co-occurring with functional HRd signatures. The misclassification of tumors with respect to a particular repair deficiency may impact clinical outcomes. The results also identified a paradoxical relationship between mutational burden and chromosomal instability. Unfortunately, patient cohorts that include comprehensive DDR profiling are small, further limiting supervised machine learning effectiveness for capturing the full DDR alteration spectrum.

To address these challenges, we developed CHIMERA-DDR (Comprehensive Heterogeneity Informing Molecular Evaluation and Risk Assessment), a probabilistic model for dynamic DDR-deficiency prediction from individual tumor samples (Fig. 4A-B, Supplementary Fig. 1). CHIMERA integrates 51 weighted molecular features encompassing DDR pathway gene mutations, functional signatures, large-scale genomic aberrations, and tumor intrinsic characteristics like ploidy and cellularity, representing a significant advancement over single-biomarker approaches (Supplementary Table 1). At its core, this framework integrates genomic aberrations, tumor intrinsic features, and genomic signatures to determine sample molecular subtypes accurately (refer Supplementary Document 2 for CHIMERA-DDR development strategy).

**Fig. 4:**
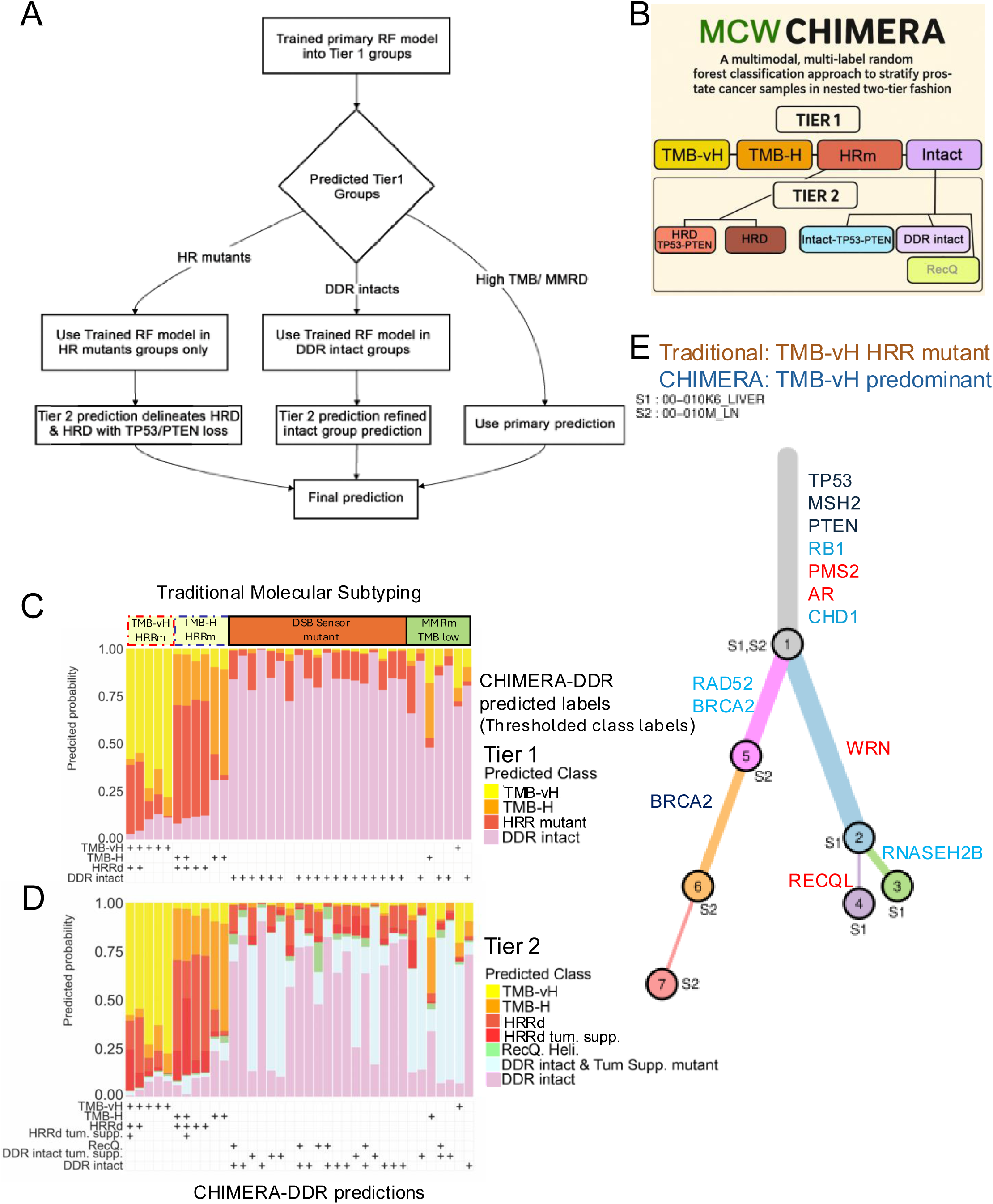
Accurate Dynamic Classification of Advanced Prostate Cancer Using a Multimodal Multilabel Random Forest-Based Nested Classification Model. **(A)** Schematic flowchart illustrating CHIMERA-DDR architecture employing a unique nested classification strategy that first delineates major 4 subclasses, then identifies more nuanced finer subclasses to infer 7 total subgroups. This model integrates 51 genomic features and signature scores from patient tumor genome and transcriptome sequencing data within a probabilistic framework to predict the relative admixture of DDR-deficiency phenotypes. **(B)** Schematic illustration of CHIMERA-DDR’s 2-tier nested classification architecture enabling hierarchical DDR subtype prediction. **(C & D)** Component barplot percent representation of CHIMERA-predicted probability decomposition for TMB-vH and TMB-H tumors with HRR mutations. Results demonstrate that TMB-vH HRR mutant subgroup tumors exhibit predominantly TMB-vH-like (MMRd-like) characteristics, whereas TMB-H HRR mutant subgroups display prevalently HRRd-like phenotypes. One of the 19 tumors with DSB sensor mutations is HRRd positive in Tier-1 analysis with Tier-2 ambiguity while defining fine-scale HRRd subgrouping, suggesting potential low-penetrance nature of these genomic events. **(E)** Clonal analysis of TMB-vH-HRR mutant sample 00-010 reveals a truncal MMRd mutation in MSH2 gene likely drives the MMRd-associated hypermutator phenotype, with minor clone S2 harboring additional potentially subclonal copy-loss events in BRCA2 and CHD1 genes. Branch length: Represents the relative mutational burden or evolutionary distance (Relative frequency of somatic events). Node width: Indicates clonal prevalence (cellular fraction that carries those somatic events).

CHIMERA-DDR employs a two-tier nested Random Forest strategy that addresses the fundamental challenge of singleton cutoff approaches across cancer subtypes by readjusting relative signature weights based on tumor intrinsic features and clonality to generate combined probability scores for DDR subclass assignment. Tier-1 stratify patients into four primary components: TMB-vH, TMB-H, HRRm, and DDR-intact phenotypes. Tier-2 provides refined subcategorization, stratifying the HRRd subset by tumor suppressor gene status (intact versus mutant TP53/PTEN) while subdividing DDR-intact groups into DDR-intact, DDR-intact with tumor suppressor mutations, DSB sensor mutants, and RecQ helicase pathway loss-of-function mutations (Fig. 4A-B). Notably, 77.59% of all HRRm tumors as per CHIMRA prediction also harbor HRRd molecular signatures and are classified as HRRd major or HRRd minor (considering mutation clonality, mutation penetrance to phenotype, or mutation zygosity, also refer Supplementary Document 1 for group definitions and Supplementary Document 2) (Supplementary Table 8A-B).

CHIMERA-DDR demonstrated technically robust classification performance across multiple validation strategies, including both cross-validation on the training dataset and independent test set validation (n=37). A core focus of the model validation was to ensure the chosen CHIMERA-DDR model was not suffering from overfitting, resulting in poor performance on the unseen real-world test dataset. In leave-one-out cross-validation (LOOCV), it achieved robust performance with the area under the curve (AUC) of 0.999(0.9969, 1.0000), 0.998(0.9957, 1.0000), 0.982(0.9685, 0.9958), 0.921(0.8552, 0.9868), 0.797(0.6693, 0.9250), 0.982(0.9684, 0.9952) and 0.976(0.9631, 0.9898) in 7 Tier-2 classes respectively. Also, consistent performance was observed in 3-fold and 5-fold cross-validation, which highlighting the model stability and absence of overfitting (Supplementary Document 2).

Notably, some molecular subgroups had limited sample sizes (e.g., RecQ helicase subgroup with n=9), precluding reliable 5-fold cross-validation for these rare subtypes. This highlights the challenge of class imbalance in molecularly heterogeneous cohorts and likely explains the more modest performance for RecQ helicase pathway prediction (AUC 0.797). While kernel density estimation (KDE)-based oversampling could have been adopted to address this imbalance, we intentionally chose to build our model exclusively on labelled real-world data to ensure clinical translatability. Despite sample size constraints, the concordance between LOOCV and 3-fold CV across all subgroups validated the model’s stability and generalizability.

Fine-scale HRRd subset deconvolution based on TP53-PTEN loss status demonstrated an AUC of 0.919 for the tumor suppressor mutant subset, indicating robust performance for stratifying patients with complex overlapping DDR phenotypes. Importantly, in every case, the model also inferred the probability of the tumor behaving as DDR-intact, a critical concept that was not considered in past models. The model showed limitations in RecQ helicase pathway prediction, achieving an AUC of 0.797, representing an area for future improvement (Fig. 5B).

**Fig. 5:**
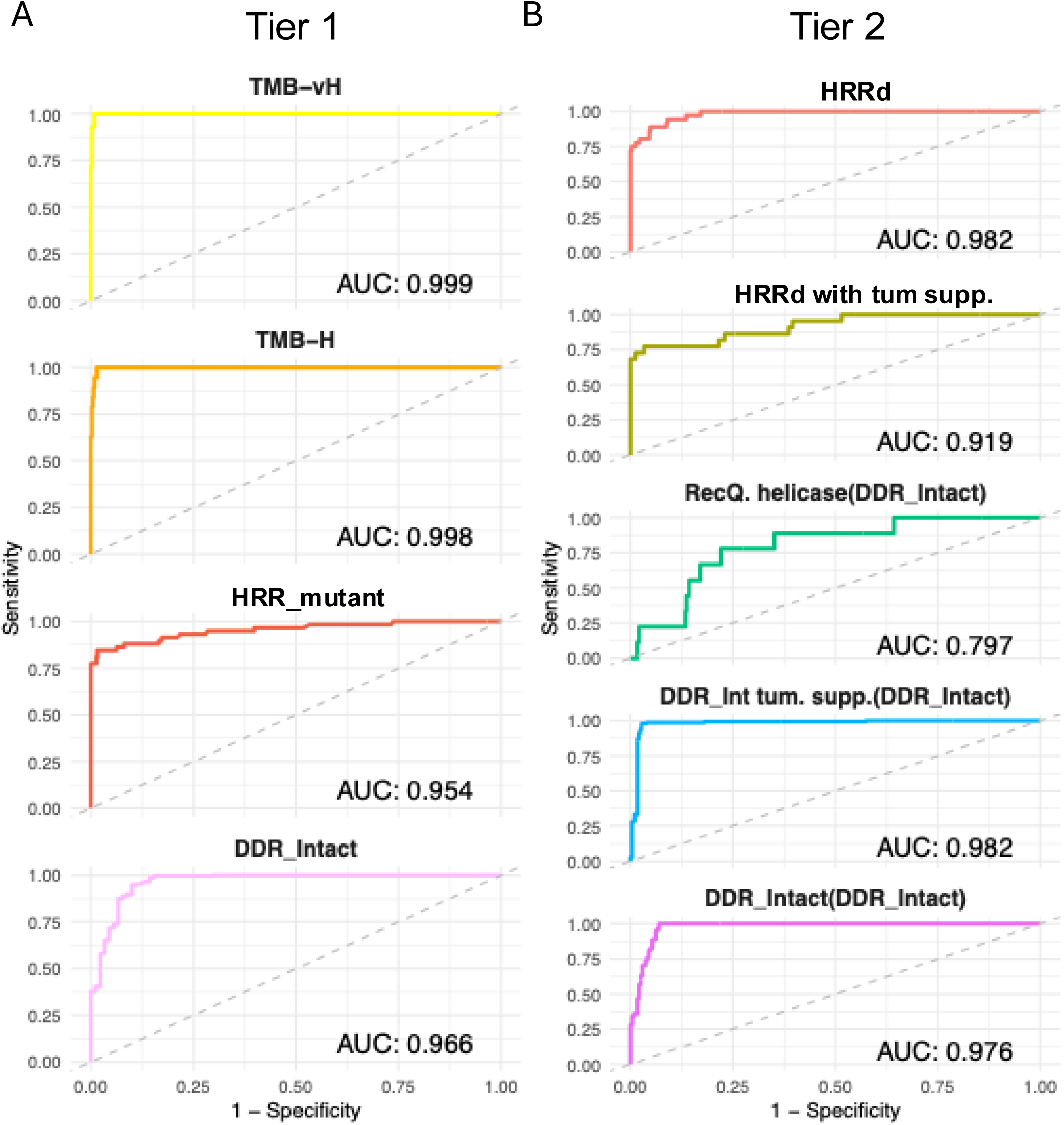
Leave-one-out cross-validation (LOO-CV) derived AUC-ROC curves showing superior technical performance in both Tier-1 and Tier-2 predictions. CHIMERA-DDR stratifies patients into 4 DDR axis molecular subgroups. Tier-2 analysis sub-stratifies the HRR mutant subgroups into HRRd with loss-of-function of selective tumor suppressor genes and those without tumor suppressor gene loss. Notably, tumor suppressor gene mutations in TP53, RB1, and PTEN impact the manifestation of HRRd-associated signature profiles and disease pathobiology. Similarly, in Tier-2 DDR-intact tumors divided into 3 molecular subgroups based on RecQ helicase pathway activity and tumor suppressor gene mutation status in the setting of DDR-intact molecular phenotype for accurate fine-scale class determination. CHIMERA-DDR demonstrated superior performance in deriving 7 molecular subgroups after Tier-2 analysis (TMB-vH, TMB-H, HRRd, HRRd tumor suppressor, RecQ helicase mutant, DDR-intact tumor suppressor, and DDR-intact without tumor suppressor mutations). Tier-1 + 2 complete analysis infers 7 fine-tuned molecular subgroups demarcated in the figure with solid blue-filled boxes in the top left of the respective AUC-ROC plot areas.

Of the 482 total tumors, 445 were used for CHIMERA-DDR model training and 37 were withheld for model test purposes, but not for model validation (5 TMB-vh HRRm (n=5), TMB-H HRRm (n=6), MMR gene mutant TMB-Low (n=7), DSB Sens. gene mutants (n=19) (Fig. 4B-D, Supplementary Table 9A). Despite the limited sample size, this unbiased approach validated CHIMERA’s ability to identify overlapping DDR phenotypes and revealed significant heterogeneity within established DDR categories.

Among the 5 TMB-vH HRRd samples, 2 harbored substantial HRRd signature components, suggesting a deeper MMRd-like phenotype with concurrent MMRd and HRRd features (Fig. 4C barplot from the left 1 and 2), while the remaining 3 (Fig. 4C barplot 3 through 5) exhibited predominantly MMRd phenotypes with HRRd appearing as a passenger event (Fig. 4C-E, Supplementary Table 9A). Similarly, within the 6 TMB-H HRRd samples, 4 demonstrated predominant HRRd phenotypes with substantial TMB-H features, while 2 samples displayed below threshold HRRd signatures, suggesting passenger mutation but predominant TMB-H characteristics, potentially indicating limited MMRd-like features (Fig. 4D-E). To clarify further, these two samples’ TMB-H characteristics are not suggestive of an MMRd-like feature. The DSB sensor subset (n=19), consisting primarily of tumors with ATM/CHEK2 mutations, showed one tumor, which is borderline HRRd negative as per the defined confidence threshold. The majority exhibited DDR-intact behavior or low-penetrance HRRd-like phenotypes, suggesting that ATM/CHEK2 mutations alone do not reliably predict HRRd status (Fig. 4D).

Testing of the 7 MMRm-low TMB subset revealed 1 tumor with HRRd-like phenotype scores despite no clear mutational explanation (Supplementary Document 1, refer to MMRm definition). This sample exhibited high LOH scores, borderline but sub-threshold HRD-Scar scores, TP53 bi-allelic loss-of-function mutations, BRCA2 monoallelic copy loss, and a focal copy-neutral LOH event in BRCA2, together indicating a complex genomic state. These features were accurately considered within CHIMERA-DDR and inferred an accurate class identity, underscoring the reliability of the model’s prediction (Fig. 4C-D). A second sample within this subset demonstrated borderline MMRd-like features with significant TMB-High characteristics despite a detected TMB of 9.8 mutations/Mb. This tumor harbored MSH2 bi-allelic loss-of-function and MLH1 loss-of-function mutations and tested positive for COSMIC MMRd signatures, potentially indicating functional MMRd despite being below the 10 mutations/Mb mutational burden threshold (47). These findings underscore the clinical utility of probabilistic modeling approaches for capturing the full spectrum of DDR pathway alterations and their complex interactions in the prostate cancer genome.

### Semi-Independent Test Set Evaluation of CHIMERA-DDR Class Determination Performance Captures Intra- and Inter-Tumor DDR Phenotype Heterogeneity

To evaluate the performance of CHIMERA-DDR, we analyzed whole exome sequences (WES) from 270 mPC tumors derived from 112 patients enrolled into a rapid postmortem tissue collection protocol (UW-TAN) (Supplementary Fig. 7A). Consistent with Kumar et al. 2015, we found limited intratumor heterogeneity in heavily treated multiple metastatic tissues from the mCRPC rapid autopsy series (Supplementary Table 10) (48). Substantial heterogeneity in TMB classification was detected in 4 of 112 patients, where tumors from the same patient included TMB-vH, TMB-H, and TMB-Low categories (Supplementary Table 10). Similarly, class switching in MSI scores was observed in tumors from 2 patients (MSI-High vs. MSI-Low). In contrast, a greater degree of intratumor heterogeneity was seen in LOH and HRD-Scar scores. To determine whether DDR axis subtypes were associated with organ-specific metastatic tropism, we analyzed site distributions; however, no statistically significant enrichment was observed, indicating that different mechanisms contributing to DDR subtypes do not preferentially drive site-specific metastasis in this dataset (Supplementary Fig. 7B), though limited sample sizes restricted robust inference (14).

To assess whether CHIMERA-DDR predictive probabilities reflect DNA from admixed normal cells in the tumor sample, we analyzed cellularity fractions in 445 training samples. For DDR-intact tumors, CHIMERA’s Tier-2 DDR-intact probability consistently exceeded the inferred normal fraction from Sequenza cellularity estimates, barring three exceptions (normal fraction = 1 - cellularity) (Fig. 6A). We investigated whether CHIMERA’s elevated DDR-intact predictions could be attributed to normal DNA contamination alone or arise from a combination of normal contamination and genuine tumor-intrinsic DDR-intact behavior (Supplementary Fig. 8A-B). Critically, DDR-deficient tumors, particularly TMB-vH and TMB-H cases, demonstrated DDR-intact prediction probabilities substantially lower than the corresponding normal DNA content (Supplementary Fig. 8B, Supplementary Fig. 9). This key observation demonstrates that if normal DNA contamination were driving DDR-intact predictions, the predicted probabilities should equal or exceed the normal fraction. Additionally, DDR component probabilities lower than tumor DNA content in DDR-deficient tumors strongly suggest that CHIMERA accurately captures the overall DDR phenotype, which is not solely driven by tumor cellularity. Of note, cellularity was included as one of the 51 input features in training CHIMERA (Feature rank #41). The cellularity estimate would assist us in inferring tumor-only DDR components.

**Fig. 6:**
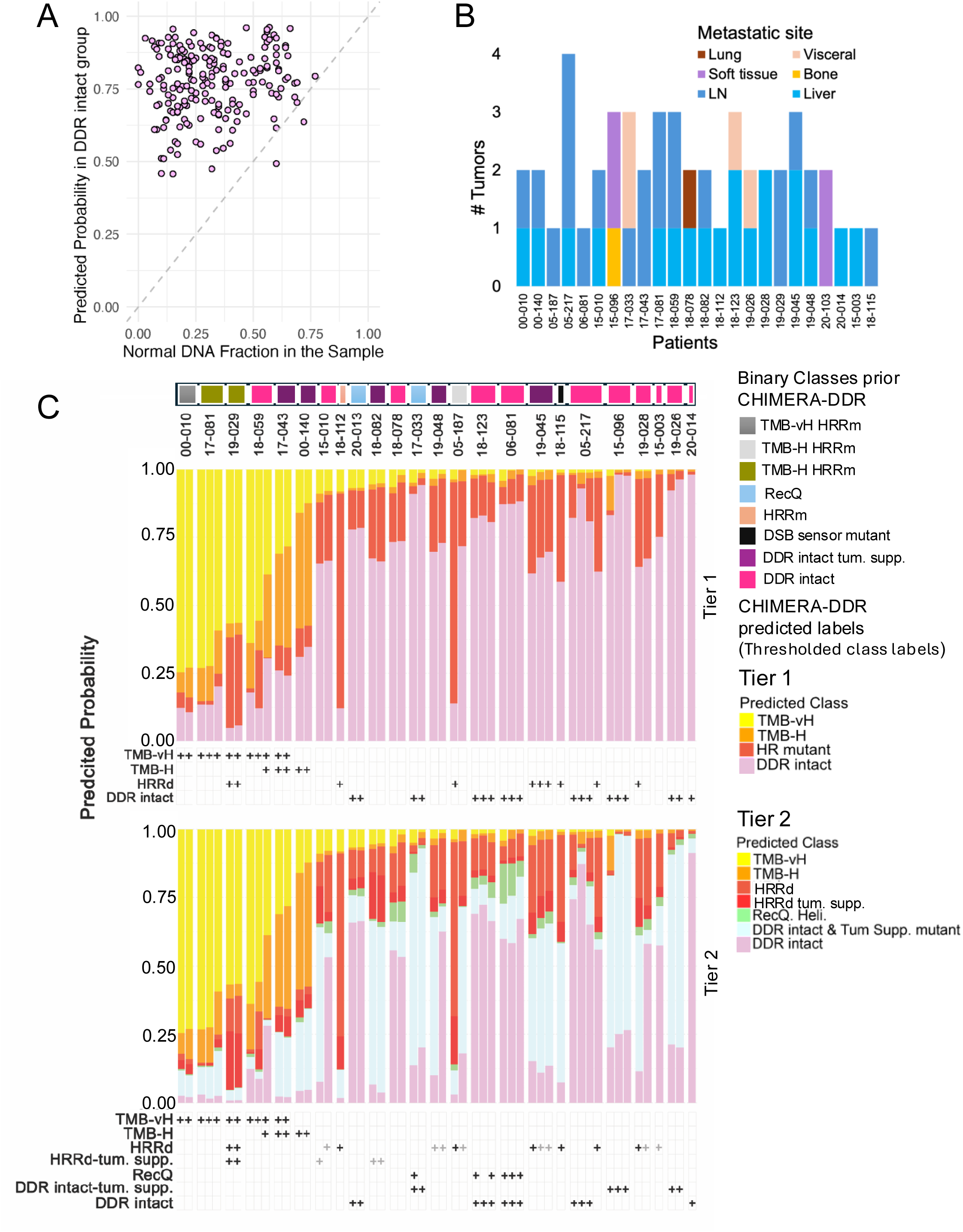
CHIMERA-DDR Demonstrates Unique Capability to Predict Admixed / Concurrent DDR-Intact Subtype Probabilities and Captures Inter-Tumoral Heterogeneity in Advanced Prostate Cancer. **(A)** Scatter plot comparing CHIMERA’s predicted DDR-intact probability versus normal DNA fraction in tumor samples. For likely DDR-intact tumors, CHIMERA’s overall Tier-2 DDR-intact prediction probability consistently exceeded the actual normal fraction inferred by Sequenza cellularity (normal fraction = 1-cellularity), initially conceived these probabilities might reflect cumulative contributions of normal DNA fraction and tumor-intrinsic DDR-intact phenotypes. Extended analysis demonstrates that DDR-intact predicted probability is not a function of sample tumor cellularity but rather is primarily guided by the tumor’s DDR-intact-like molecular characteristics, tumor’s clonal structure indicating tumor-intrinsic DDR functional states detection; also refer Supplementary Fig. 7-9. **(B)** Distribution of 52 tumor samples from 24 patients across multiple metastatic sites (lung, visceral, soft tissue, lymph node, bone, liver) from the UW-TAN case series covering 5 DDR subgroups with corresponding whole genome sequencing data. This test series enabled evaluation of CHIMERA’s robustness for DDR status inference using signature scores derived from multiple metastatic sites. **(C)** Stacked bar plots showing Tier-1 and Tier-2 DDR subtype prediction probabilities across multiple tumor samples from the same patients. Each bar represents relative prediction probabilities for DDR subtypes (TMB-vH, TMB-H, HRR mutant, DDR-intact) in Tier-1, with refined subcategorization in Tier-2 (DSB sensor/TP53/PTEN mutations, BRCA2 mutations, PTEN/TP53 mutations, and no mutations). Asterisks indicate tumors with CHIMERA-DDR intratumoral heterogeneity. Analysis reveals substantial homogeneity in DDR-deficiency prediction probabilities within patient groups, with notable heterogeneity observed in select cases.

Among the 58 HRR-mutant tumors in the training set, CHIMERA predicted that 77.59% (n=45) exhibited HRRd-like molecular characteristics. In contrast, the remaining 22.41% showed HRRd-like integrative genomic features below the determined threshold (Supplementary Table 8A & Supplementary Table 8B). Detailed curation identified contributing factors, including subclonal loss-of-function events, hypomorphic alleles, and lower-penetrance HRR aberrations that diminished the functional impact on DDR phenotype. Integrative analysis further revealed tumors designated as HRRd that unexpectedly displayed DDR-intact mutational signatures. Together, these findings reconcile the apparent paradox and highlight CHIMERA-DDR’s ability to capture the full functional spectrum of DDR pathway dysfunction, moving beyond rigid binary models toward probabilistic resolution of overlapping and heterogeneous phenotypes.

To evaluate CHIMERA’s robustness for DDR status inference, we selected 24 patients (52 tissue genomes) from the UW-TAN case series covering 5 DDR subgroups with corresponding whole genome sequencing (WGS) data. (Fig. 6B, Supplementary Table 9B) noted a limited degree of discordance in TMB estimates between WES and WGS, potentially reflecting underlying heterogeneity in mutation burden across coding and noncoding regions (Supplementary Table 10). Notably, both datasets were analyzed using similar pipelines with the same quality cutoffs. However, due to the consistently lower depth of coverage in WGS, low allele frequency somatic variants were often missed. Conversely, WGS demonstrated superior sensitivity for detecting structural variants, which was not detected in WES analysis. This unique paired dataset enabled evaluation of CHIMERA’s prediction accuracy and ability to capture inter-tumoral heterogeneity while inferring admixed DDR phenotype probabilities. We performed 2-tier DDR subtype prediction and plotted Tier-1 probabilities in percent component bar plots to understand intratumor versus inter-tumor heterogeneity, with each bar plot component representing relative prediction probability (Fig. 4E, 6C, Supplementary Fig. 10; also refer to Supplementary Document 3). We investigated whether multiple tumor samples from the same patient clustered based on DDR-deficiency components. WGS-derived input data revealed substantial intra-patient homogeneity in deconvolved DDR-deficiency probabilities. Four patients had only a single tumor sample with WGS data available, precluding their inclusion in this analysis. Of the remaining 20, three patients exhibited substantial heterogeneity, predominantly in HRRd and DDR-intact/DDR-intact tumours. supp. probability components, whereas the other seventeen demonstrated homogeneous profiles across multiple samples.

## DISCUSSION

This study fundamentally reframes DNA damage repair classification in advanced prostate cancer by demonstrating that conventional biomarker approaches inadequately capture therapeutically relevant biology. We observed that tumors with concurrent DDR pathway alterations exhibit variable therapeutic responses to immune checkpoint inhibitors versus PARP inhibitors despite carrying conventional biomarkers predictive for both. Through comprehensive analysis of 672 advanced PC exomes, we present three transformative findings that challenge current precision medicine paradigms in DDR-targeted therapy.

### TMB-Very High as a Distinct Molecular Entity

We established that TMB-very high tumors (≥18.61 mut/Mb) represent a molecularly distinct entity from conventional TMB-High tumors, with profound clinical implications for immunotherapy selection. This data-driven threshold emerged from systematic analysis demonstrating that higher TMB cutoffs achieve superior accuracy in identifying MMRd phenotypes, revealing a fundamental specificity-sensitivity trade-off critical for therapeutic decision-making (49–55). Importantly, MMR pathway gene bi-allelic mutants with paradoxically low TMB (<10 mut/Mb) demonstrate that mutation-based detection strategies are insufficient; these tumors fail to exhibit functional MMRd characteristics despite definitive genetic alterations (Bi-allelic loss of MSH2, MSH3).

TMB-very high tumors exhibit MMR-driven biology with preserved genomic integrity, robust CD8+ T-cell infiltration, and PD-L1 upregulation, while paradoxically maintaining AR-driven transcriptional programs. This “inflamed but AR-active” phenotype contradicts prevailing models of AR independence in hypermutated PCs and suggests these tumors may require different therapeutic approaches than conventional TMB-High cases. While conventional TMB ≥10 mut/Mb cutoffs show limited predictive value in PC, our elevated threshold demonstrates improved correlation with immunotherapy response, consistent with observations that patients with TMB above median levels show enhanced checkpoint inhibitor responses in metastatic castration-resistant PC (23,24). We statistically inferred the TMB-vH subgroup to enrich for definitive MMRd-like tumors, then performed deep molecular characterization to develop our dynamic classification model. This genotype-molecular admixture complexity reflects biological reality and underscores why probabilistic approaches like CHIMERA are essential for precision DDR stratification.

### Functional DDR Pathway Dominance in Molecularly Complex Tumors

We discovered that tumors harboring both HRRd and MMRd alterations do not exhibit dual therapeutic vulnerabilities as predicted by conventional biomarkers. Instead, these molecularly complex tumors display functional dominance of one DDR pathway over another, explaining why patients respond to either PARPi or immunotherapy, but not necessarily both. CHIMERA-DDR reveals this biological reality by quantifying relative DDR pathway contributions within individual tumors rather than imposing binary classifications, capturing tumor heterogeneity and functional interrelationship (epistatic) between HRR and MMR pathways. Critically, 24% of HRR-mutant tumors displayed MMRd-predominant phenotypes rather than expected HRRd biology. For example, tumor 00-010 carried both TMB-vH and HRR mutations, yet CHIMERA correctly identified its dominant MMRd phenotype, a finding confirmed by clonality analysis showing major MSH2 loss with only minor BRCA2 alterations (Fig. 4E, Supplementary Fig. 10).

### Superior DDR-deficiency Classification Beyond Genotype-Only Approaches

CHIMERA-DDR represents a paradigm shift from genotype-only HRRd classification to functional phenotype prediction, addressing the critical gap between genomic alterations and therapeutic vulnerability. Our analysis demonstrates that 77.59% of HRRm tumors exhibited HRRd-like molecular characteristics while 22.41% did not, revealing substantial genotype-phenotype discordance missed by conventional approaches. CHIMERA-DDR identifies TMB-vH and TMB-vH HRRm as clinically distinct molecular subgroups in prostate cancer with unique therapeutic vulnerabilities. CHIMERA’s ability to predict DDR-intact-like phenotypes alongside DDR-deficient signatures adds crucial clinical insight, particularly for detecting DDR-intact clones that may require combination targeted approaches.

CHIMERA’s biological validation confirmed molecular characteristic inference through multiple lines of evidence. Tumor cellularity analysis revealed that DDR-deficient tumors had substantially lower DDR-intact prediction probabilities compared to normal DNA content (1-cellularity), indicating that CHIMERA probabilities represent weighted molecular characteristics of the tumor composition at the bulk tumor level (Fig. 6A, Supplementary Fig. 11A-B). Semi-independent validation using paired whole genome sequencing data from 52 samples across 24 patients demonstrated substantial intragroup homogeneity in predicted probabilities across metastatic sites, confirming stable, biologically meaningful DDR phenotypes (Fig. 6C).

### Clinical Implications for Combination Therapy Strategies

CHIMERA-DDR’s findings may offer insights into why PARPi+ICI combination strategies have shown limited success in prostate cancer. Phase 3 KEYLYNK-010 trial (pembrolizumab + olaparib) was stopped for futility (11,12,47,53–56), while phase 2 studies including CheckMate 9KD (nivolumab + rucaparib) and JAVELIN PARP Medley (avelumab + talazoparib) demonstrated only modest activity, primarily in biomarker-selected subgroups. These trials enrolled molecularly heterogeneous populations where genomically similar tumors may have exhibited fundamentally different functional molecular types, potentially diluting treatment effects. Our discovery that concurrent DDR alterations display pathway dominance rather than dual vulnerabilities suggests even tumors with multiple alterations may respond optimally to pathway-specific monotherapy rather than combination approaches.

CHIMERA represents a fundamental shift from binary classification toward dynamic probabilistic inference that captures the biological reality of admixed DDR phenotypes in cancer. By integrating multiple molecular features, CHIMERA better approximates functional DDR status beyond simple mutation presence, enabling accurate stratification of patients across the full spectrum of DDR alterations. Critically, CHIMERA-DDR infers clonal versus subclonal architecture of DDR mutations, a distinction overlooked by conventional approaches. Even bona-fide MMRd or HRRd tumors often retain degrees of DDR-intact-like behavior; CHIMERA quantifies this residual functional capacity, enabling clinicians to match therapeutic strategy to clonal complexity. This is particularly valuable for tumors with TMB-High and concurrent HRR aberrations, where clinicians currently face difficult choices between immunotherapy and PARPi treatments due to conflicting molecular signals. CHIMERA resolves this uncertainty by separating mixed DDR signatures and identifying mutation-positive tumors that actually behave like DDR-intact cancers, providing accurate classification for both HRR and MMR pathways— capabilities impossible with current single-cutoff methods. Tumors with mixed clonal architecture may require combination or sequential therapies, while clonally uniform deficiencies warrant pathway-specific monotherapy. Such integrated molecular profiling may better identify patients who would benefit from ICI monotherapy versus combination approaches, directly addressing the current clinical challenge of therapeutic selection in molecularly complex DDR-altered tumors.

CHIMERA-DDR’s utility will expand alongside emerging therapies. New treatments, including CTLA-4 and PD-L1 combinations, PD-L1-targeted CAR-T cells, and next-generation DNA-damaging agents, will require similar sophisticated patient selection. This framework extends beyond prostate cancer to any malignancy where DNA repair deficiencies guide treatment decisions. However, prospective clinical trials with treatment outcomes are essential to validate whether CHIMERA-predicted phenotypes accurately identify responders to immunotherapies or DNA-damaging therapies, such as PARP inhibitors.

## METHODS

### Study Cohorts

We assembled a comprehensive multi-modal genomic dataset from advanced PC patients across multiple independent cohorts. Fundamentally, our framework was designed to integrate genomic variants with genomic signatures for precise molecular subtype classification of individual samples. The primary WES discovery cohort (n=672) comprised samples from two independent sources: the University of Washington rapid autopsy program (UW-TAN, n=270, Supplementary Table 1) and Stand Up To Cancer Dream Team consortium (SU2C, n=444). After quality control filtering to remove low-coverage sequence data, 466 high-quality samples were retained for downstream analysis. RNA sequencing data (n=656) were obtained from overlapping patients across both cohorts: SU2C samples (n=482) sequenced in two batches (n=270 and n=212) and UW-TAN samples (n=174) sequenced as a single batch (Supplementary Table 1). For clinical validation, we assembled an independent cohort of 130 patients with clinical-grade genomic sequencing results from four academic medical centres (Supplementary Table 7). This study was approved by the Institutional Review Board under protocols PRO00044894 and PRO00053119. All patient data were appropriately deidentified prior to analysis to ensure confidentiality and compliance with ethical standards. To assess intra-tumoral heterogeneity, we analyzed a test cohort (n=52) comprising multiple metastatic tumor sites with paired whole-genome sequencing data from 24 patients, generating 51 CHIMERA-DDR input features to evaluate model performance across diverse sampling contexts (Supplementary Table 9A-B,10).

### DNA Sequence Analysis

WES and WGS reads were aligned to the human reference genome (GRCh38/hg38) using Burrows-Wheeler Aligner-MEM v0.7.17.1 (57). GATK Best Practices workflows were implemented for preprocessing, including alignment quality assessment, read sorting with Picard MarkDuplicates (v2.24.1), and Base Quality Score Recalibration (BQSR v4.1.9.0) to ensure high-quality variant calling (58).

### Variant Calling, Annotation & Tumor Mutation Burden estimation

Germline variants were called using GATK HaplotypeCaller with population database filtering (gnomAD v3.1.1, 1000 Genomes, ESP6500) to exclude common variants (allele frequency >0.1%) (59). Somatic variants were identified using MuTect2 and Strelka v2 in tumor-normal paired mode, requiring minimum coverage of 12 high-quality reads (Q≥40) with at least 6 supporting alternate alleles (60,61). A union set of filtered mutations from both callers maximized detection sensitivity. BEDTools filtration retained variants within target capture regions for accurate TMB inference. All variants underwent manual curation, and TMB was calculated using pyTMB (v1.4.0), incorporating the union variant set, effective genome size, and cellularity (62).

### Microsatellite Instability Analysis

Microsatellite instability (MSI) status was determined using MSIsensor-pro v1.2.0 from paired tumor-normal sequencing data, with the same analytical pipeline applied to both whole-exome and whole-genome sequencing datasets (63). MSI-High (MSI-H) tumors were defined using an MSI score threshold ≥3.5, while microsatellite-stable (MSS) tumors had MSI scores <3.5, following established clinical guidelines for PC. This classification enabled stratification of tumors based on MMRd status for downstream DDR pathway analysis.

### Pathogenicity Classification

Somatic pathogenic mutations in 46 DNA repair pathway genes and associated tumor suppressors were identified through comprehensive annotation using ANNOVAR and validated with Mutalyzer 2.0. The curated gene panel encompassed homologous recombination repair (BRCA1, BRCA2, PALB2, RAD51B, RAD51C, RAD51D), DSB sensors (MRE11A, NBN, ATM, CHEK1, CHEK2), mismatch repair (MLH1, MSH2, MSH6, PMS2), Fanconi anemia pathway (FANCA, FANCB, FANCC, FANCD2, FANCE, FANCF, FANCG, FANCI, FANCL, FANCM), RecQ helicases (BLM, WRN, RECQL4, RECQL5, RECQL), and DNA repair-associated genes (TP53, PTEN, RB1, CDK12, RNASEH2B, ATR, AR, SPOP)(36,37,61,64–74). Germline pathogenic variants were annotated following ACMG recommendations (75), while somatic pathogenic mutations were classified based on: (1) nonsense mutations, (2) frameshift insertions/deletions, (3) canonical splice site alterations (±1,2 bp), (4) missense mutations with established functional impact, and (5) previously reported pathogenic variants in clinical databases. Copy number alterations were assessed using Sequenza CNV pipeline. Gene-specific copy number aberrations were performed using a homebrew copy number gene annotation pipeline using pyTHiA-CNA (Supplementary Document 5). Bi-allelic inactivation is defined as two concurrent hits involving pathogenic mutations, complex structural aberrations, and/or copy loss. Classifications were cross-validated against cBioPortal, ExAC, Kaviar, ClinGen, ClinVar, ClinVitae, CIViC, OncoKB, and UniProt databases (69,74,76–79). Identical analytical pipelines and quality control thresholds were applied to whole-genome sequencing analysis.

### Mutational Signature Analysis

Cosmic mutational signatures were identified using SigProfilerExtractor (v1.1.23) with the COSMIC v3.4 reference signature set (80). Analysis employed non-negative matrix factorization (NMF) with 100 replicates for statistical robustness, resampling enabled for stability assessment, and 10,000-1,000,000 iterations with convergence tolerance of 1e-15 using GRCh38 reference and opportunity genomes with default trinucleotide context. Signatures of clinical interest included HRRd (C. signature 3), MMRd (signatures 6, 14, 15, 20, 21, 26, 44), and DNA polymerase epsilon deficiency (POLE; signatures 9, 10a-d, 14, 20).

### Copy Number Analysis, LOH & HRD-Scar

Copy number analysis utilized paired tumor and normal depth of coverage and B-allele frequency data. Copy number variations, tumor cellularity, and purity were determined using Sequenza v2.0.0 (78). Tumor ploidy and purity were derived alongside segmented allele-specific copy numbers from allele-specific copy number analysis. Copy number annotation was performed using a homebrew annotator tool referencing GENCODE v45 annotation. Loss of heterozygosity (LOH) scores were calculated from copy number calls using the method described by Abkevich et al. (81). We counted toward this score events that were greater than 15 Mb in length and defined by a non-zero copy number count with an inferred minor allele count of 0 (81). Chromosomes that were affected by such LOH events over ≥75% of their entire length were excluded from both numerator and denominator, as they typically arise through non-HRRd associated mechanisms. HRRd scores were computed as the sum of LOH, telomeric allelic imbalance (TAI), and large-scale state transitions (LST) using the HRD-Scar tool by Sztupinszky et al (82). Bi-allelic inactivation was defined as concurrent pathogenic mutation and copy number loss (log2 ratio <-0.5 and B-allele frequency indicating loss of heterozygosity) as well as complex structural aberrations.

### Subgroup-Specific Consensus Genomic Aberration Analysis

To generate subgroup-specific consensus profiles of genomic aberrations, we established a CNVkit-based analytical framework for comparative copy number analysis across molecular subgroups (83). Discovery cohort samples were stratified into DDR-defined molecular subgroups, and individual BAM files within each subgroup were pooled to create representative datasets. Pooled BAM files were downsampled to 300× depth of coverage using Picard tools to normalize sequencing depth and reduce computational burden while maintaining copy number detection sensitivity (84). CNVkit (v0.9.12) was then applied to the downsampled pooled BAM files to generate consensus copy number profiles, enabling identification of subgroup-specific recurrent genomic alterations and comparative analysis of copy number landscapes across different DDR molecular subtypes (83).

### Transcriptome Analysis

RNA sequencing reads were aligned to the hg38 reference genome using STAR v2.7.3a, with gene-level expression quantified using GenomicAlignment Bioconductor package (85). Differential gene expression analysis was performed using edgeR and limma packages with filterByExpr filtering and Benjamin-Hochberg FDR correction (86,87). The RNA-Seq dataset (n=656) comprised SU2C (n=482) and UW-TAN (n=174) cohorts. Only poly(A) capture samples were analyzed to avoid batch effects from different library preparation methods, confirmed by multidimensional scaling analysis (41,43,88).

The 31-gene cell cycle progression (CCP), 10-gene androgen receptor activity(ARa), and 10-gene neuroendocrine activity (NEa) scores were computed using GSVA on log₂ FPKM RNA-Seq data with validated gene panels (41,43). All scores were standardized across the cohort for comparative analysis.

### Immune Infiltration Analysis

Immune cell infiltration analysis was performed independently on each cohort using CIBERSORT v1.04 with the LM22 signature matrix to estimate the relative abundance of 22 immune cell subsets from RNA-Seq data (88). Analysis was conducted with absolute mode enabled and statistical significance evaluated using 100 permutations per cohort. Following individual cohort analysis, mean and median infiltrating immune cell frequencies were calculated for each DDR molecular subgroup to enable comparative assessment of immune microenvironment profiles across different DNA repair deficiency states.

### Clonality and Tumor Evolution Analysis

Tumor clonal evolution and phylogenetic reconstruction were performed using PhyloWGS v1.0-rc2 to infer evolutionary histories from multi-sample tumor data (89). Input preparation utilized the PhyloWGS parser to integrate high-quality VCF variant calls, segmented copy number alterations from Sequenza, and tumor cellularity estimates. Simple somatic mutations in DNA repair pathway genes were prioritized during phylogenetic reconstruction to enhance evolutionary accuracy and focus on DDR-relevant clonal dynamics. PhyloWGS analysis employed Markov Chain Monte Carlo (MCMC) sampling with four independent parallel chains to ensure convergence and robust posterior distribution sampling. Chain convergence was assessed using standard MCMC diagnostics, and posterior probabilities were calculated for phylogenetic tree topologies and clonal frequency estimates. The resulting phylogenetic trees and clonal evolution patterns were visualized using Clonevol v0.1, which generated comprehensive plots displaying subclonal architecture, clonal frequency dynamics across multiple tumor sites, and evolutionary trajectories (90). This analysis enabled assessment of clonal heterogeneity patterns across different DDR molecular subtypes and identification of founder versus acquired DDR alterations during tumor progression.

## AUTHORS’ CONTRIBUTIONS

K. Sharma: Data curation, formal analysis, validation, investigation, visualization, methodology, writing–original draft, project administration, writing–review and editing. D.A. Wilson: data curation, software, formal analysis, visualization, methodology, writing–original draft, writing–review and editing. Y. Wang: software, formal analysis, visualization, methodology, writing– original draft, writing–review and editing. S. Haidar: formal analysis, methodology, investigation, data curation. V. Bhatlapenumarthi: formal analysis, methodology, investigation, data curation. D. Boiarsky: formal analysis, investigation, data curation. J. M. Cipriaso: formal analysis, investigation, Data curation. L. Yermakov: formal analysis, investigation, data curation. R. Dutta: formal analysis, methodology. M. Vashisth: review and editing, I. M. Coleman: formal analysis. A. Bankhead: formal analysis. Shantanu K. Jadhav: data curation, resources, investigation. B. Neupane: formal analysis, resources, investigation, data curation. B. W. Taylor: resources, writing–review and editing. S. Rao: review and editing, M. T. Nevalainen: review and editing, A. A. Nelson: writing–review and editing. C. Morrissey: review and editing, M. T. Schweitzer: writing–review and editing. R. B. Montgomery: writing– review and editing. E. S. Antonarakis: Writing–review and editing. P.G.P: Writing–review and editing. J. E. Berchuk: resources. K. Zarrabi, resources, writing–review, and editing. C. C. Pritchard: resources, writing–review and editing, G. Ha: resources, writing–review and editing, A. Kothari: resources, writing– review and editing, B. Jeorge: review and editing, H. Chen: review and editing, R. Kurzrock: writing– review and editing, P. S. Nelson: resources, writing–review and editing, P. L. Auer: formal analysis, resources, writing–review and editing, A. Banerjee: formal analysis, resources, writing–review and editing. D. Kilari Conceptualization, resources, data curation, supervision, writing–critical review and editing. N. D. Sarkar: Conceptualization, resources, data curation, software, formal analysis, supervision, funding acquisition, validation, investigation, visualization, methodology, writing–original draft, project administration, writing–review and editing.

## ACKNOWLEDGMENTS

We extend special thanks to William Branson and Matthew Flister from the MCW Research Computing Core, and Jason Erb from MCW IS for facilitating the setup of computational terminals, platforms, and environments, and for their consistent support throughout this data-intensive research. We thank the MCW Biostatistics, Geospatial, Epidemiology and Outcomes Shared Resources Core members. We thank the patients for their families for their altruistic contributions to this study. We deeply thank participating patients & families, Evan Yu, Heather Cheng, Jessica Hawley, Daniel Lin, Funda Vakar-Lopez, Michael Haffner, Martine Roudier, Lawrence True, Meagan Chambers, Peter S. Nelson, and the rapid autopsy teams for their contributions to the University of Washington Prostate Cancer Donor Rapid Autopsy Program. We thank Amit Joshi, Todd Miller, Gustavo Leone, and Steven Kroft for their consistent support and encouragement throughout the successful execution of this research. We particularly thank Neeraj Agarwal for critically evaluation of our preliminary results. No generative artificial intelligence tools were utilized for data analysis, figure generation, or computational code development. Large language model assistance was employed in a limited capacity exclusively for language editing and manuscript polishing (subscribed tools from anthropic, OpenAI).

## Data Availability

The SU2C whole-exome sequencing data and SU2C bulk RNAseq data are publicly available at: https://github.com/cBioPortal/datahub/tree/master/public/prad_su2c_2019.

The UW-TAN whole-exome sequencing data publicly available at: https://github.com/cBioPortal/datahub/tree/master/public/prad_fhcrc

UW-TAN Bulk RNA-Seq data: Gene Expression Omnibus repository (GEO) IDs: **GSE147250**, **GSE171729**, **GSE228283**.

The Whole genome sequence data used in this study are generated as the part of UW-TAN cohort Whole genome sequencing effort. Tertiary analysis files: mutations (VCF) and copy number analysis (*.segment.txt) files are available at: https://github.com/Niel20/CHIMERA_DDR_Validation_Cohort_52_WGS.

## Code Availability

The computer code associated with the CHIMERA-DDR including the trial data and scripts can be accessed at (https://github.com/Niel20/CHIMERA), all rights are reserved to Copyright 2025 Medical College of Wisconsin. For academic usage codes would be made available by request to corresponding author at.

