## Supplementary Document 1 for "CHIMERA-DDR: A Machine Learning Framework for Classifying Heterogeneous Mismatch-Repair and Homologous-Recombination Deficiency Patterns in Prostate Cancer"

**Defining 11 molecular classes of advanced prostate cancers along the DNA repair axis**

Tumor mutation burden is very high with concurrent HRR mutation (>18.61 mut/Mb , TMB-vH_HRRm): Samples with concurrent very high tumor mutation burden (≥18.61 mut/mb) and loss-of-function genomic aberrations in the homologous recombination repair pathway genes.

Tumor mutation burden high with concurrent HRR mutation (≥10-18.61 mut/Mb, TMB-H_HRRm): Samples with concurrent high tumor mutation burden (≥10 mut/mb) and loss of function genomic aberration in the homologous recombination repair pathway genes.

TMB-very High (TMB-vH): Tumors with a tumor mutational burden ≥18.61 mut/Mb.
 These tumors fall into the hypermutated category, reflecting extensive genomic instability and higher predicted immunogenicity.

TMB-High (TMB-H): Tumors with a mutation burden greater than 10 mutations per megabase (mut/Mb) and less than 18.61mut/mb are classified as TMB-high.

MMRm TMB-Low: Tumors with biallelic mismatch repair deficiency (loss-of-function mutations/LOH in MMR genes) but with low TMB (<10 mutations/Mb).

Homologous Recombination pathway gene mutant (HRRm): Homologous Recombination Repair mutant (HRRm) refers to a cellular state in which the homologous recombination (HR) DNA repair pathway is impaired or non-functional. In this study, HRRm was defined by the presence of pathogenic germline or somatic loss-of-function mutations in HR pathway genes, with or without a second hit in the same gene.

Homologous Recombination pathway gene mutant with Tumor Suppressor Gene bi-allelic loss of function mutation (HRRm tumor suppressor or HRRm Tum. Supp.): A subset of HRRm tumors that also carry biallelic loss of tumor suppressors (TP53, RB1, PTEN) in addition to biallelic HRR mutations.

Double Strand Break Sensor gene mutants (DSB Sens.): Tumors with biallelic inactivation of genes responsible for DNA double-strand break sensing and signaling. DSB sensor gene list included ATM, CHEK2, MRE11, NBN, DNA2 and EXO1.

RecQ-Helicase Gene Mutant (RecQ-Heli): Tumors with biallelic loss-of-function alterations in RecQ helicase genes (WRN, BLM, RECQL5, RECQL, RECQL4).

DDR-intact with Tumor Suppressor Gene bi-allelic loss of function mutation (DDR-Int. Tum. Supp.): Tumor-Suppressor Gene mutant: Tumors without DDR pathway defects but with biallelic loss of tumor suppressors (*TP53, RB1, PTEN*).

DDR-intact: Tumors with no biallelic alterations in the DDR pathways gene (HRR, MMR, DSB sensing, RecQ helicases) or selected tumor suppressors.

**Defining 7 molecular decomposition classes of advanced prostate cancer along the DNA repair axis inferred by CHIMERA-DDR**

CHIMERA-DDR is a probabilistic framework developed to stratify patients into seven molecular subgroups based on 51 curated genomic features and mutational signature scores.

**TMB-very High (TMB-vH):** Tumors in this category demonstrate molecular behavior closely resembling that of functionally MMR-deficient tumors, reflecting hypermutation and profound genomic instability. TMB-High (TMB-H): Compared with the TMB-vH subgroup, these tumors display a lower degree of hypermutation, indicating increased but not extreme genomic instability. While not reaching the profound hypermutated phenotype of TMB-vH tumors, the TMB-H group still represents a distinct molecular state associated with intermediate immunogenicity and potential therapeutic relevance.

**Homologous Recombination Pathway Based Repair deficient (HRRd):** Tumors with a predicted HRRd post CHIMERA tier 2 analysis, based on probabilistic integration of HRR-related genomic features and HRRd-associated mutational signatures. These tumors exhibit patterns consistent with homologous recombination repair deficiency.

**Homologous Recombination pathway with HRRd and Tumor Suppressor Gene bi-allelic loss of function mutation (HRRd-Tum. Supp):** A refined subclass within HRRd, predicted by CHIMERA-DDR tier 2 analysis, where tumors not only demonstrate HRRd-associated genomic signatures but also show strong probability scores for biallelic loss of tumor suppressor genes (TP53, RB1, PTEN), indicating compounded genomic instability.

**RecQ-Helicase Gene Mutant (RecQ-Heli):** Identified by CHIMERA-DDR as tumors with genomic feature patterns and mutational signatures consistent with loss of RecQ helicase function (e.g., BLM, WRN, RECQL family), representing a distinct mechanism of DDR dysfunction.

**DDR-intact with Tumor Suppressor Gene bi-allelic loss of function mutation (DDR Int. Tum. Supp.):** Predicted by CHIMERA-DDR as DDR proficient tumors that nonetheless display biallelic inactivation of tumor suppressors (TP53, RB1, PTEN), suggesting oncogenesis is driven by checkpoint/tumor suppressor loss rather than canonical DDR failure.

**DDR-intact:** tumors with a probability distribution strongly favoring the intact DDR state, showing no significant HRRd, MMRd, DSB-sensor, or RecQ-associated signature enrichment or genomic aberrations. These serve as the reference “baseline” molecular subgroup.
