## Supplementary Document 2 for "CHIMERA-DDR: A Machine Learning Framework for Classifying Heterogeneous Mismatch-Repair and Homologous-Recombination Deficiency Patterns in Prostate Cancer"

CHIMERA-DDR (Classifier for Heterogeneous Integrated Mutational Events in DNA Repair) is a machine learning–based framework designed to delineate complex DNA damage repair (DDR) deficiency states in prostate cancer. Unlike traditional binary approaches that classify tumors solely as HRRd (homologous recombination deficient) or DDR-intact, CHIMERA-DDR integrates multi-omic features—including genomic scars, transcriptomic profiles, MSI, genomic mutations and pathway-level signatures into an 7 class nested random forest model.

This multimodal stratification enables the identification of overlapping or concurrent DDR-deficiency states. Improved prediction of therapeutic response to PARP inhibitors, immune checkpoint blockade, and emerging precision medicine strategies. Enhanced biological interpretation of heterogeneous DDR landscapes across clinical cohorts. By capturing the dynamic and multifaceted nature of DDR-deficiencies, CHIMERA-DDR provides a more nuanced framework to advance biomarker discovery and therapeutic decision-making in advanced prostate cancer.

CHIMERA-DDR model development (https://github.com/Niel20/CHIMERA): We adopted a hierarchical classification approach motivated by the inherent nature of DNA repair deficiency-associated genomic aberrations, recognizing that DNA repair defects exist on a molecular spectrum rather than as discrete categories. This biological reality reflects the complex interplay of subclonal mutation patterns, variable penetrance of detected mutations to phenotype, and the differential importance of pathway genes in defining molecular subtypes. Such heterogeneity necessitates nested analysis to capture the underlying mechanisms driving different mutational processes and their resulting fine-scale molecular classifications.

Feature Ranking: The CHIMERA-DDR random forest classifier identifies the relative importance of multiple DDR features through Gini importance metrics. Estimated feature importance rankings are outlined in Supplementary Fig. 11A-B, Supplementary Fig. 12A. We noted Tier-1 feature rankings are substantially different from those of Tier-2 rankings.

More precisely, we customized the random forest framework to accommodate the nested group structure present in the training dataset. Initially, a primary random forest model was trained to classify observations into Tier-1 categories: TMB-vH, TMB-H, HRR mutant, and DDR-intact (see Supplementary Document 1 for definitions). Subsequently, for the Tier-1 groups TMB-H and HRR mutant, we trained two additional, secondary random forest models to further classify these groups into their respective Tier-2 subcategories HRRd and HRRd with tumor suppressor gene loss-of-function (HRRd tum. Supp.). This two-stage approach results in a hierarchical random forest architecture, as illustrated in the figure below, enabling refined prediction while preserving the underlying group hierarchy. Notably, the Tier-1 probability cut-off for HRRm and DDR-intact is considered the gateway criterion for evaluating Tier-2 probability. The final inference regarding the fine-tuned groups is based directly on the Tier-2 evaluation.

Model Cross Validation (CV): We adopted leave-one-out cross-validation (LOOCV), considering variability within training subgroups and limited sample sizes for some subgroups. The iterative process ensures maximum training data usage in each iteration, maintains the unbiased prediction on the leave-one subject. Using the aggregated predictions from the LOOCV procedure, the model performance is evaluated by AUC scores along with their confidence interval in each subgroup, which represent the model's ability to distinguish closely related molecular subtypes—critical for clinical applications where misclassification between biologically similar groups could impact therapeutic decision-making. In particular, DeLong method (DeLong et al. (1988)) is used to evaluate the uncertainty of an AUC. Meanwhile, the similar procedure was done in 3-fold and 5-fold cross validation, where we pick one fold as the testing dataset, the rest folds was serve as the training dataset. Iterating all the folds so that every fold can be the testing dataset. In the end, the similar AUC score was achieved, ensuring the stability of CHIMERA-DDR.

**Leave One Out Cross Validation: AUC (CI)**

*** Tier 1**

Group TMB-vH AUC: 0.999(0.9969, 1.0000)

Group TMB-H AUC: 0.998(0.9957, 1.0000)

Group HR_Mutant AUC: 0.955(0.9197, 0.9894)

Group DDR_Intact AUC: 0.967(0.9438, 0.9899)

*** Tier 2**

Group TMB-vH AUC: 0.999(0.9969, 1.0000)

Group TMB-H AUC: 0.998(0.9957, 1.0000)

Group HR mutant AUC: 0.982(0.9685, 0.9958)

Group HRD tum. supp. AUC: 0.921(0.8552, 0.9868)

Group RecQ. helicase AUC: 0.797(0.6693, 0.9250)

Group DDR_Int tum. supp. AUC: 0.982(0.9684, 0.9952)

Group DDR_Intact AUC: 0.976(0.9631, 0.9898)

**3-fold Cross Validation: AUC (CI)**

*** Tier 1**

Group TMB-vH AUC: 0.999(0.9981, 1.0000)

Group TMB-H AUC: 0.996(0.9913, 1.0000)

Group HR_Mutant AUC: 0.958(0.9223, 0.9929)

Group DDR_Intact AUC: 0.966(0.9411, 0.9915)

*** Tier 2**

Group TMB-vH AUC: 0.999(0.9981, 1.0000)

Group TMB-H AUC: 0.996(0.9913, 1.0000)

Group HR mutant AUC: 0.982(0.9666, 0.9973)

Group HRD tum. supp. AUC: 0.926(0.8571, 0.9940)

Group RecQ. helicase AUC: 0.836(0.7280, 0.9448)

Group DDR_Int tum. supp. AUC: 0.983(0.9703, 0.9950)

Group DDR_Intact AUC: 0.979(0.9663, 0.9910)

**5- fold Cross Validation: AUC (CI)**

*** Tier 1**

Group TMB-vH AUC: 0.999(0.9976, 1.0000)

Group TMB-H AUC: 0.996(0.9903, 1.0000)

Group HR_Mutant AUC: 0.939(0.8929, 0.9851)

Group DDR_Intact AUC: 0.956(0.9274, 0.9853)

*** Tier 2**

Group TMB-vH AUC: 0.999(0.9976, 1.0000)

Group TMB-H AUC: 0.996(0.9903, 1.0000)

Group HR mutant AUC: 0.982(0.9666, 0.9973)

Group HRD tum. supp. AUC: 0.888(0.7851, 0.9903)

Group RecQ. helicase AUC: 0.838(0.7068, 0.9696)

Group DDR_Int tum. supp. AUC: 0.980(0.9662, 0.9944)

Group DDR_Intact AUC: 0.977(0.9642, 0.9901)

Cutoff Determination: The proposed model outputs the predicted probabilities for each Tier-2 group, which correspond to the proportion of decision trees in the random forest assigning an observation to each group. To generate a confident group classification, we determined cutoff probabilities using Youden’s J statistic (Youden, 1950), which optimizes the trade-off between sensitivity and specificity. A prediction is considered confident only if the predicted probability exceeds the corresponding cutoff value.

Justification for Excluding Other Models (e.g., GLMnet, SVM): Due to missing data within the cohort, only 361 out of 445 observations have complete covariate information (Missing data on AR activity, NE activity and CCP scores). Models such as GLMnet and support vector machines (SVM) require complete-case data, limiting their applicability and resulting in reduced sample size for training. In contrast, random forest algorithms are capable of handling missing values intrinsically, thereby utilizing the full dataset. Additionally, modelling interactions between covariates in methods like GLMnet and SVM typically requires extensive manual feature engineering. In contrast, random forests naturally account for complex interactions during the tree-building process, offering a more efficient and robust approach in high-dimensional settings.

Cutoff Application: Because the proposed model is organized into two tiers, cutoffs are applied hierarchically when assigning Tier-2 classes. For instance, classification into the Tier 2 category HRRd tum. supp. requires that its probability exceed the Tier-2 cutoff (0.1844), but this condition is evaluated only if the corresponding Tier-1 cutoff (0.3175) is also satisfied.
