## Supplementary Document 3 for "CHIMERA-DDR: A Machine Learning Framework for Classifying Heterogeneous Mismatch-Repair and Homologous-Recombination Deficiency Patterns in Prostate Cancer"

Clinical validation cohort source- 130 tumors’ clinical summary and molecular evaluation occurred through cumulative collection of deidentified patients’ data evaluation. The study is supported by IRB#

i. Department of Laboratory Medicine, University of Washington
ii. Yale New Haven Hospital
iii. Medical College of Wisconsin
iv. Sidney Kimmel Comprehensive Cancer Center
