## Supplementary Document 4 for "CHIMERA-DDR: A Machine Learning Framework for Classifying Heterogeneous Mismatch-Repair and Homologous-Recombination Deficiency Patterns in Prostate Cancer"

**Statistical methods:**

Logistic Regression-based Determination of TMB Cutoff that Distinguishes MSI High Subset Efficiently: We assessed the TMB-MSI relationship by dichotomizing MSI at 3.5 (MSI > 3.5 vs. ≤ 3.5) and fitting a logistic regression model with TMB as predictor, yielding AIC 145.97 with positive coefficients indicating higher TMB associated with MSI-high status. ROC analysis identified an optimal TMB cutoff of 19.13 by minimizing Euclidean distance to the top-left corner, balancing sensitivity and specificity better than KDE-derived estimates. 3-fold cross-validation across identified an optimal cutoff at TMB 18.61 based on peak mean AUC performance, with 5-fold cross-validation selected as optimal for balancing computational efficiency with robust performance estimates across all data partitions.

Other Statistical Analysis: Enrichment analyses were conducted in R (v4.3.2) using the packages stats, dplyr, ggplot2, ggpubr, and rstatix. Chi-square tests (2×2 contingency tables) were applied to evaluate enrichment of MSI instability and MMRd status within the TMB-vH subgroup when all expected cell counts were ≥5; otherwise, Fisher’s exact test was used. More generally, Fisher’s exact test was applied for categorical comparisons with small sample sizes, and the Wilcoxon rank-sum test was used for continuous variables that did not satisfy normality assumptions (e.g., TMB, ploidy, PGA, LOH). Continuous variables (TMB, ploidy, PGA, LOH) were compared using the Wilcoxon rank-sum test due to non-normal distributions, while categorical features (MSI status, HRD-Scar) were assessed using Fisher’s exact test. Spearman’s rank correlation was performed to evaluate monotonic relationships between continuous variables, with correlation strength reported as Spearman’s rho (ρ). Correlation analyses were carried out using rstatix, and results were visualized with ggplot2 and ggpubr. All tests were two-sided, and a p < 0.05 was considered statistically significant.

Survival Analysis- Survival outcomes were evaluated using Kaplan–Meier analysis in R (v4.3.2) with the survival and survminer packages. Kaplan–Meier curves were generated to estimate survival distributions across groups, and differences were assessed using the log-rank test. All reported p-values were two-sided, with p < 0.05 considered statistically significant. Waterfall plots were generated using Prostate-Specific Antigen (PSA) data to illustrate individual patient-level responses. Patients were ordered by maximum percent change in PSA within their respective molecularly stratified groups, and bars were color-coded to highlight subgroup-specific response patterns.
