## Supplementary Document 5 for "CHIMERA-DDR: A Machine Learning Framework for Classifying Heterogeneous Mismatch-Repair and Homologous-Recombination Deficiency Patterns in Prostate Cancer"

**Supplementary Document 5:- pyTHiA-CNA - Python Toolkit for Informed Copy Number Annotation**

pyTHiA-CNA is a Python tool (version 3.7+) designed for annotating copy number segments with gene information from GTF files, requiring pandas as the primary dependency. The tool processes tab-delimited segment files and standard GTF annotation files to identify genes overlapping with copy number segments. The algorithm categorizes gene-segment overlaps into three coverage types: full coverage (fcov) when genes are completely within segments, partial coverage (pcov) for genes partially overlapping segments, and uncovered (ucov) for genes not covered by any segment. Output includes gene annotations with corresponding segment information and coverage type classification. The tool processes approximately 1000 genes per second and is optimized for WGS, WES, and targeted sequencing data. It is available under GNU GPL v3.0 at <https://github.com/Niel20/pyTHiA-CNA>.
