## Supplementary Document 6 for "CHIMERA-DDR: A Machine Learning Framework for Classifying Heterogeneous Mismatch-Repair and Homologous-Recombination Deficiency Patterns in Prostate Cancer"

**Supplementary Document 6:- PGA - Percent Genome Alteration Estimator**

The PGA Estimator is an R-based tool (version 3.6+) utilizing parallel processing for calculating Percent Genome Alteration scores from copy number variation data. The tool requires dplyr, data.table, and doParallel packages. Input consists of space-separated copy number call files derived from WGS or WES data, containing chromosome coordinates and copy numbers for both alleles. The algorithm figures out PGA scores that show how much of the genome has changes in copy numbers, giving separate scores for allele A, allele B, and a combined score for each sample. The tool implements multi-core processing for efficient batch analysis of large sample cohorts. It is distributed under GNU GPL v3.0 at <https://github.com/Niel20/PGA_Estimator>.
