## Supplementary Document 7 for "CHIMERA-DDR: A Machine Learning Framework for Classifying Heterogeneous Mismatch-Repair and Homologous-Recombination Deficiency Patterns in Prostate Cancer"

**Supplementary Document 7: - D-AV-Plotter - Dual-Allele Visualization Plotter**

D-AV-Plotter is a Python-based tool (version 3.7+) for creating multi-panel genomic visualizations that integrate bar plots, categorical annotations, and dual-allele gene mutation data. The tool requires the NumPy, pandas, Seaborn, and matplotlib libraries. It employs a dual-allele representation system where each gene is visualized through two columns (Gene_A and Gene_B), rendering homozygous states as solid rectangles and heterozygous states as diagonally split triangles with distinct colors for each allele. Input consists of two tab-separated files: a main data file containing sample information, quantitative measurements, categorical annotations, and paired gene mutation columns, and a metadata file defining plot structure and color schemes. Output generates publication-ready figures with vertically stacked panels displaying bar charts, categorical heatmaps, and gene mutation visualizations with automatic legend generation. The tool is available under GNU GPL v3.0 at <https://github.com/Niel20/D-AV-Plotter-Genomic-Data-Visualization-Tool>.
