## Supplementary figures and images for "CHIMERA-DDR: A Machine Learning Framework for Classifying Heterogeneous Mismatch-Repair and Homologous-Recombination Deficiency Patterns in Prostate Cancer"

### Supplementary Fig. 1

Supplementary Figure 1

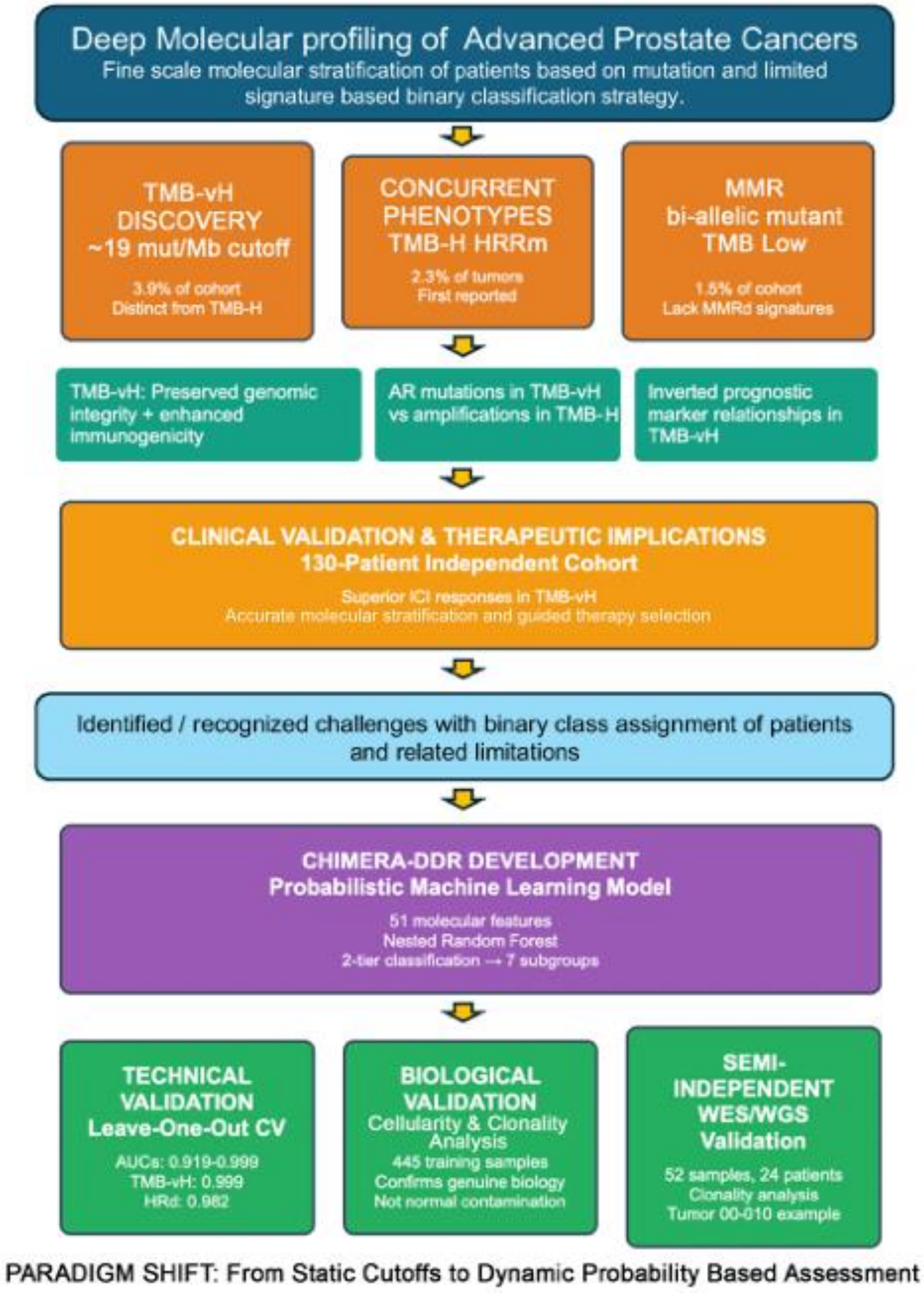

### Supplementary Fig. 4

#### Supplementary Figure 4

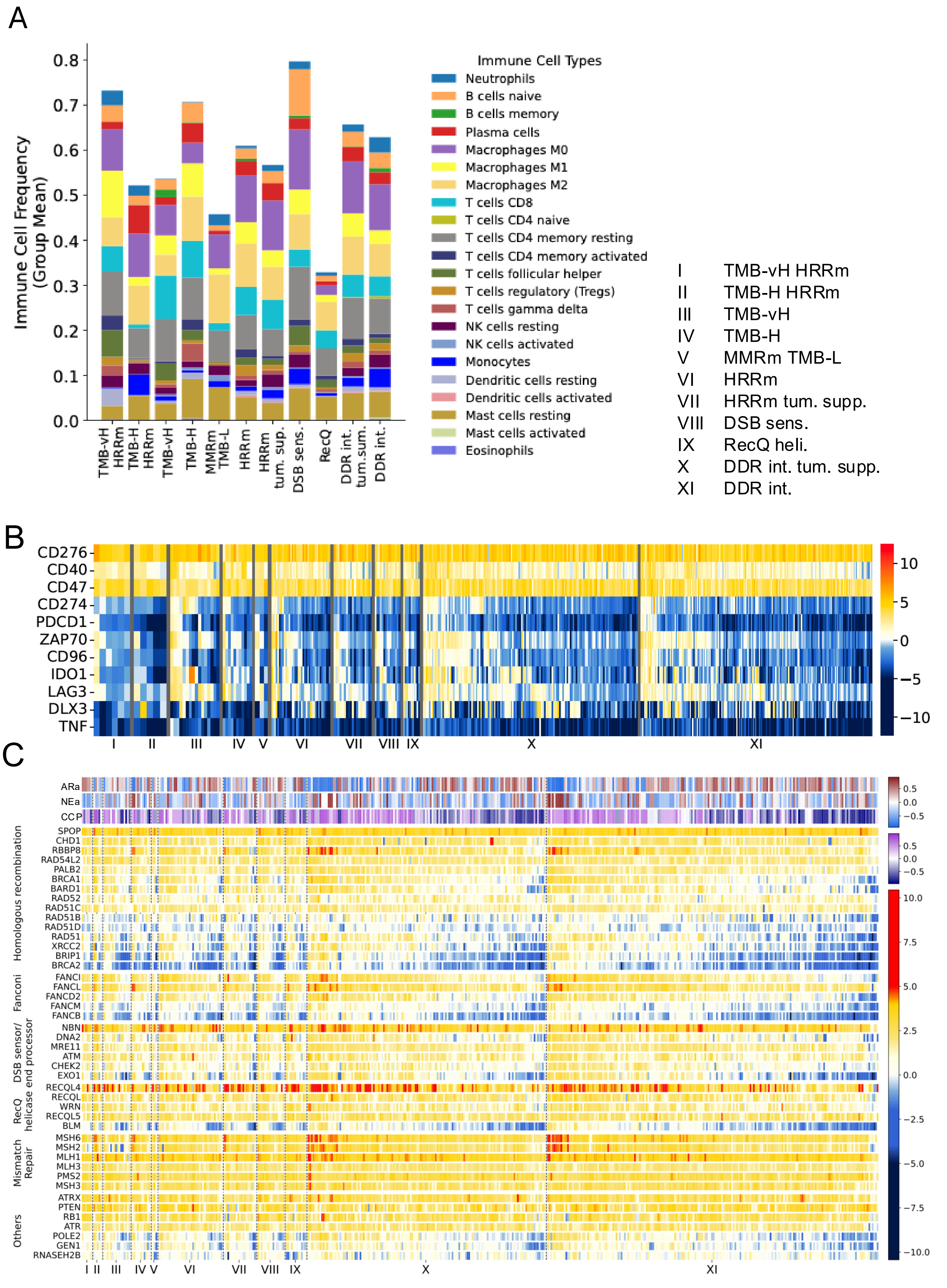

### Supplementary Fig. 7

#### Supplementary Figure 7

A

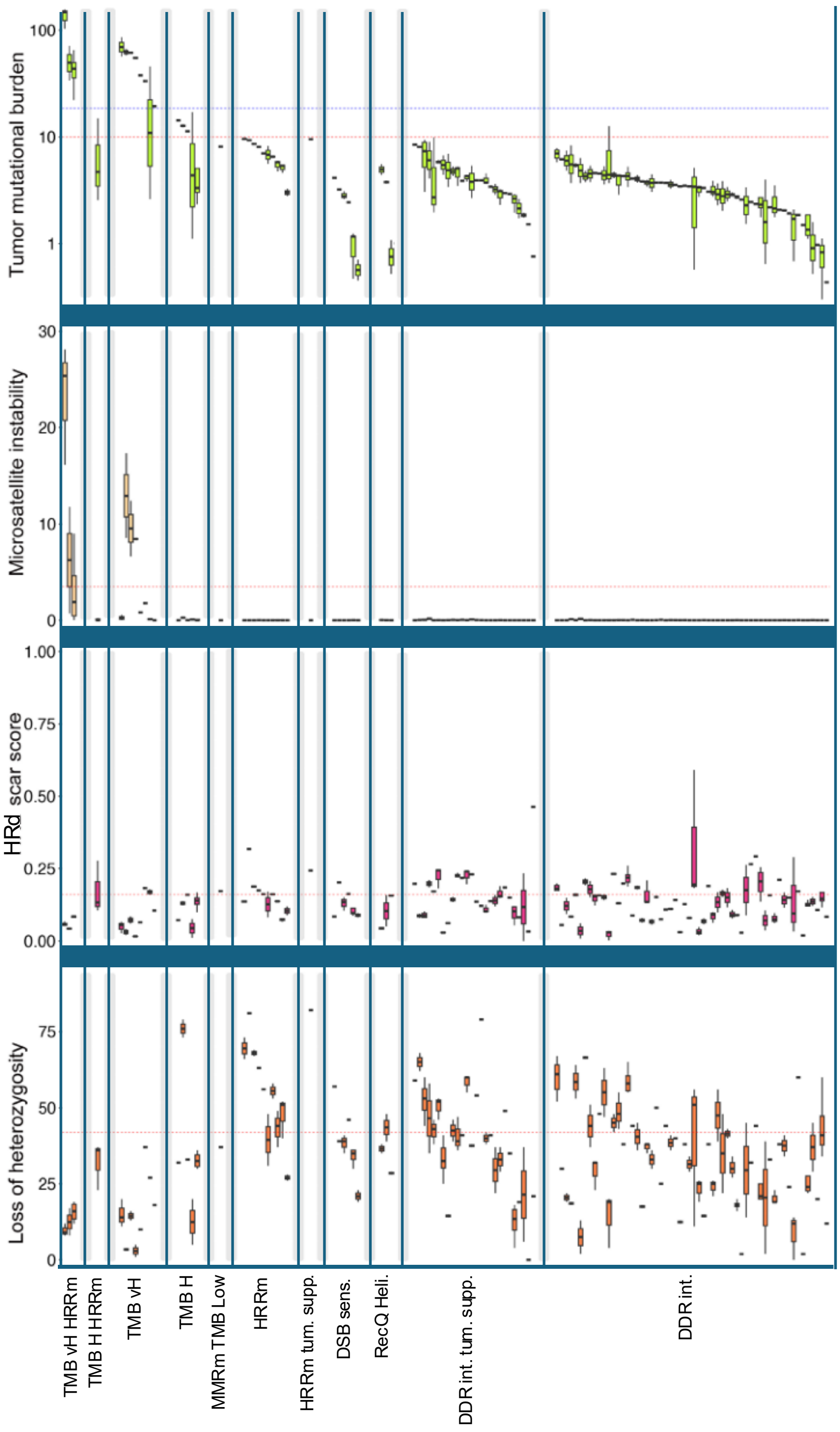


B

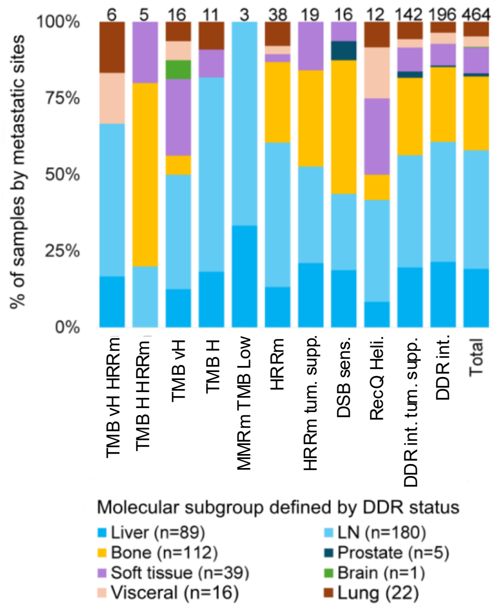

### Supplementary Fig. 8

##### Supplementary Figure 8

A

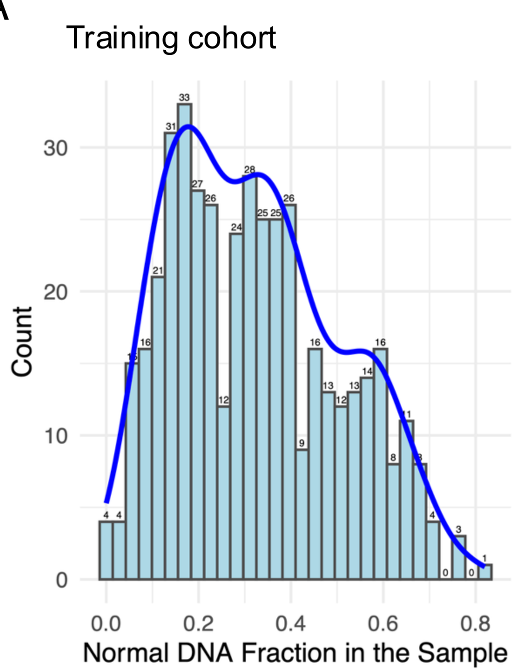


B

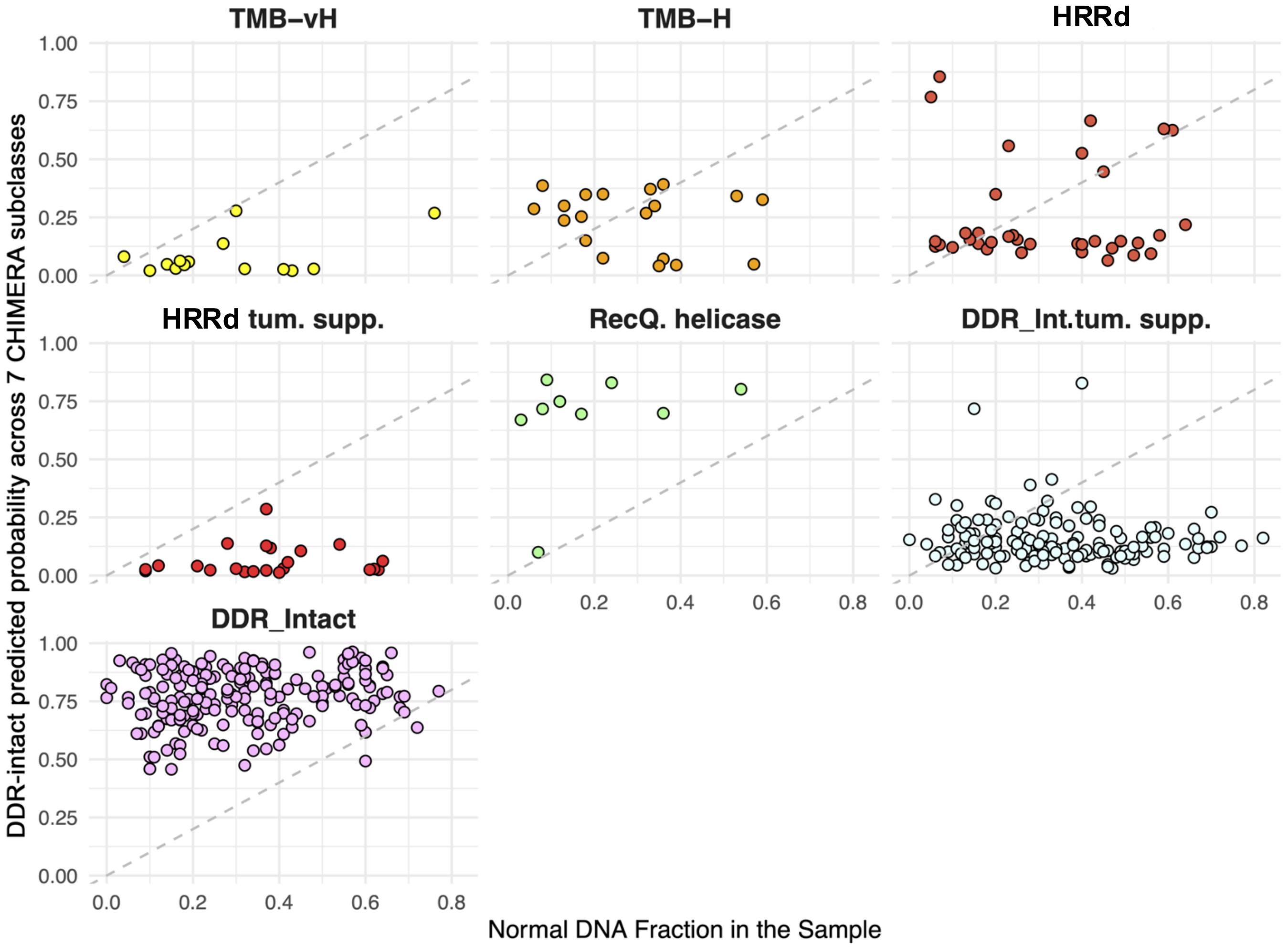

### Supplementary Fig. 9

#### Supplementary Figure 9

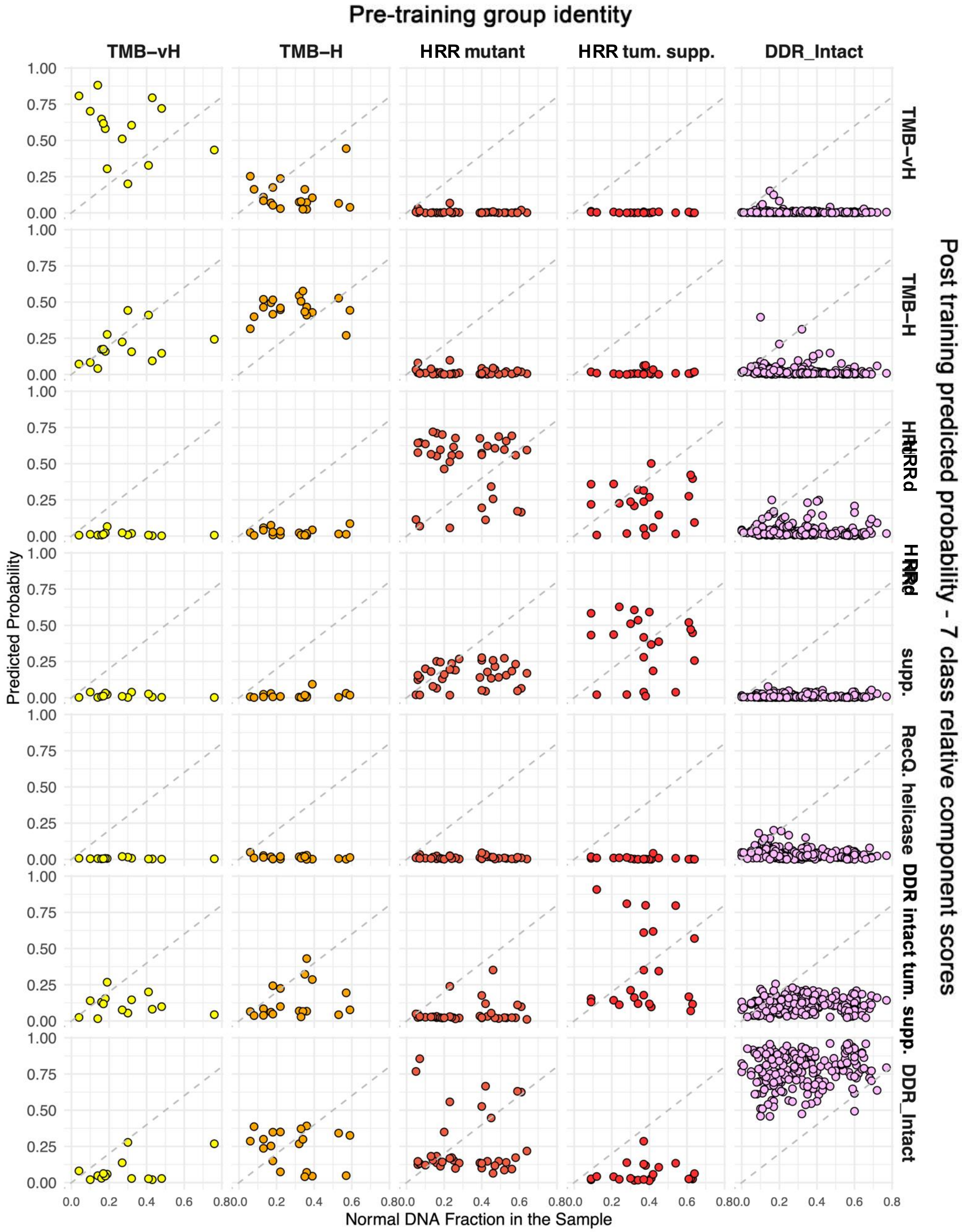
