## Supplementary Fig. 2 for "CHIMERA-DDR: A Machine Learning Framework for Classifying Heterogeneous Mismatch-Repair and Homologous-Recombination Deficiency Patterns in Prostate Cancer"

### Supplementary Figure 2

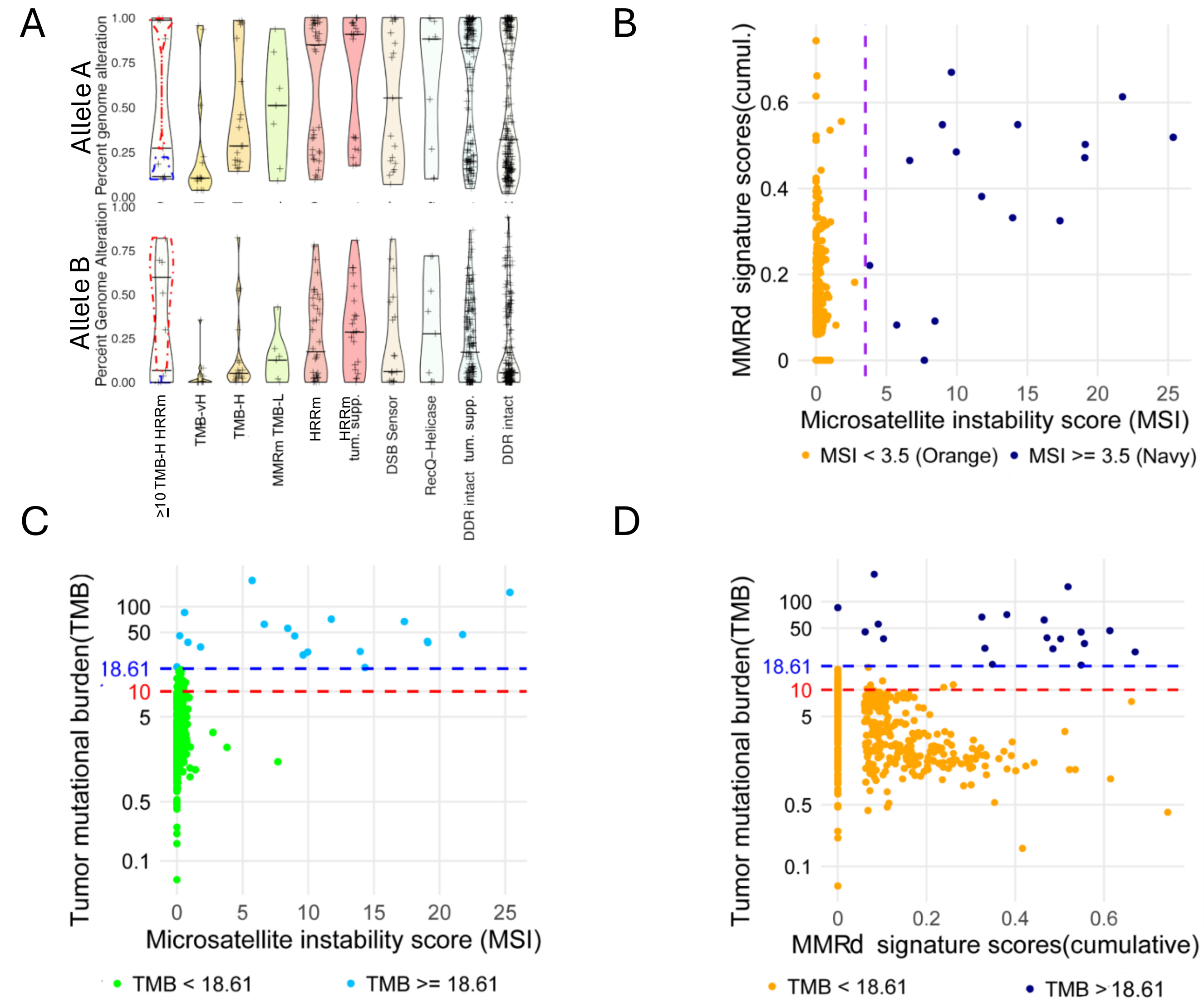


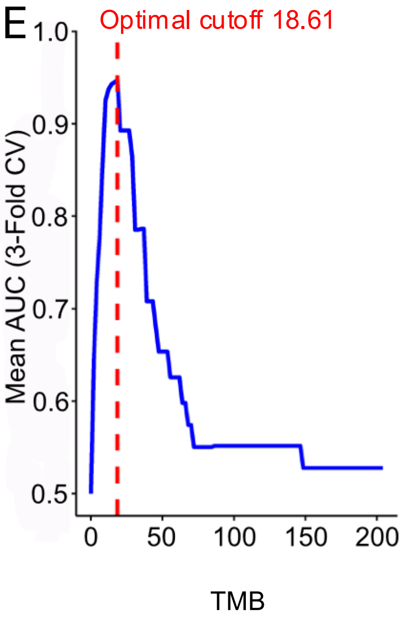


F

| MSH2 | p_value | median_group | median_intact | direction | adj_p_value |
|---|---|---|---|---|---|
| TMB-vH HRRm | 0.002757727 | 0.4297955 | 2.5899615 | Down | **0.013788633** |
| TMB-H HRRm | 0.930257891 | 2.592314 | 2.5899615 | Up | 0.930257891 |
| TMB-vH | 1.21E-05 | 1.0053825 | 2.5899615 | Down | **0.000121002** |
| TMB-H | 0.523664232 | 2.902356 | 2.5899615 | Up | 0.92435707 |
| MMRD-TMB-L | 0.354941848 | 2.577007 | 2.5899615 | Down | 0.887354621 |
| HRRm | 0.647049949 | 2.582791 | 2.5899615 | Down | 0.92435707 |
| HRRm tum. supp. | 0.752322723 | 2.572414 | 2.5899615 | Down | 0.930257891 |
| DSB Sensor | 0.894812159 | 2.607305 | 2.5899615 | Up | 0.930257891 |
| RecQ Helicase | 0.326624436 | 2.7742155 | 2.5899615 | Up | 0.887354621 |
| DDR intact tum. supp. | 0.565641062 | 2.585735 | 2.5899615 | Down | 0.92435707 |

G

| TP53 | Biallelic loss | Non biallelic | Total samples | Odds_Ratio | P_Value |
|---|---|---|---|---|---|
| TMB-vH HRRm | 0 | 5 | 5 | 0 | 0.587721628 |
| TMB-H HRRm | 1 | 5 | 6 | 0.758 | 1 |
| TMB-vH | 3 | 11 | 14 | 1.034 | 1 |
| TMB-H | 3 | 16 | 19 | 0.711 | 0.774603251 |
| MMRD-TMB-L | 3 | 4 | 7 | 2.844 | 0.169323792 |
| HRRm | 16 | 42 | 58 | 1.444 | 0.301869213 |
| DSB Sensor | 0 | 19 | 19 | 0 | 0.018374687 |
| RecQ Helicase | 1 | 8 | 9 | 0.474 | 0.691903534 |
| Intact | 72 | 273 | 345 | | |
| Total | 99 | 383 | 482 | | |
