## Supplementary Fig. 3 for "CHIMERA-DDR: A Machine Learning Framework for Classifying Heterogeneous Mismatch-Repair and Homologous-Recombination Deficiency Patterns in Prostate Cancer"

### Supplementary Figure 3

A

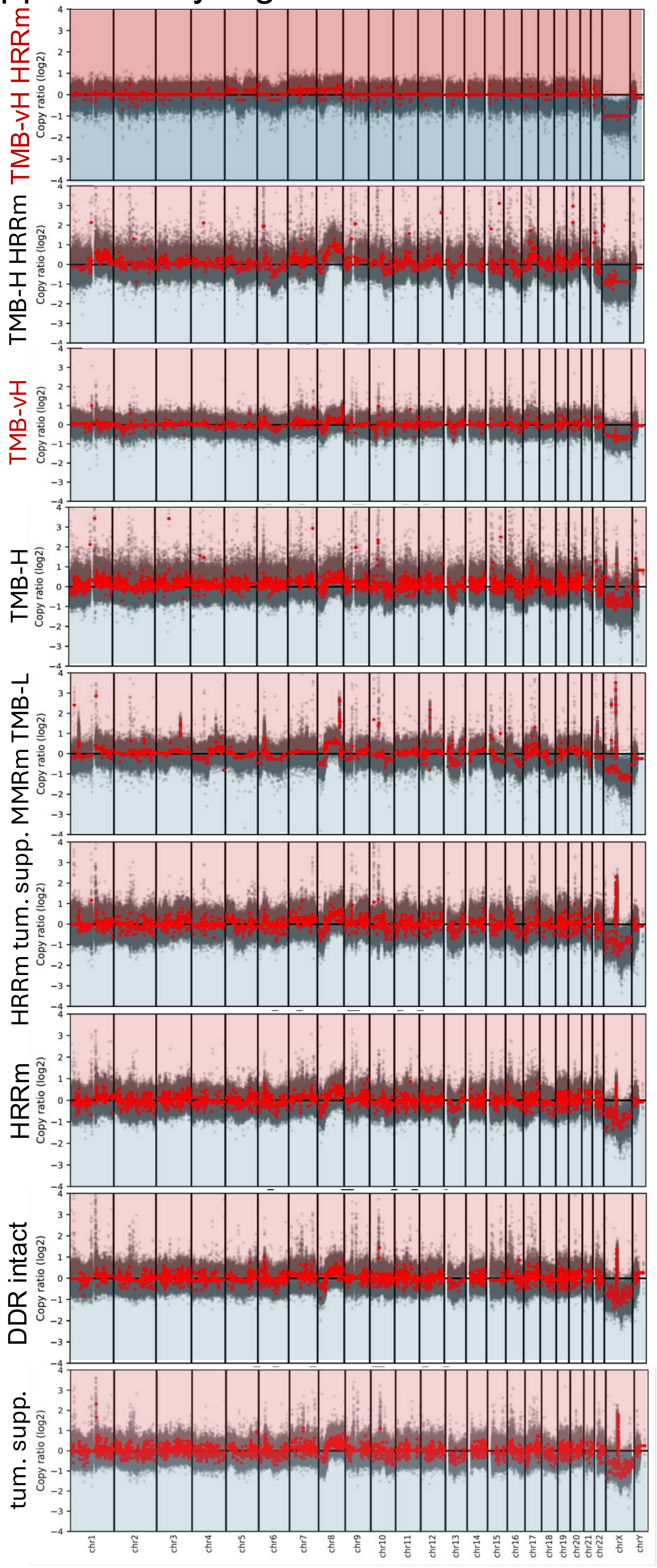


B

| Molecular subgroup | 8p loss | 8q gain | 10q loss | 13q loss | 17p loss |
|---|---|---|---|---|---|
| | % of the subgroup | | | | |
| TMB-vH HRRm | 40 | 80 | 0 | 20 | 0 |
| TMB-H HRRm | 66.7 | 100.0 | 33.3 | 33.3 | 83.3 |
| TMB-vH | 28.6 | 28.6 | 7.1 | 42.9 | 28.6 |
| TMB-H | 47.4 | 52.6 | 47.4 | 36.8 | 42.1 |
| MMRm-TMB-L | 57.1 | 57.1 | 42.9 | 42.9 | 28.6 |
| HRRm | 66.7 | 63.9 | 27.8 | 44.4 | 44.4 |
| HRRm Tum supp. | 77.3 | 63.6 | 40.9 | 63.6 | 63.6 |
| DSB Sensor | 68.4 | 63.2 | 10.5 | 36.8 | 15.8 |
| RecQ Helicase | 66.7 | 88.9 | 22.2 | 33.3 | 44.4 |
| Intact tum supp | 65.3 | 60.4 | 31.3 | 49.3 | 45.8 |
| Intact | 56.7 | 43.8 | 19.4 | 38.8 | 23.9 |
