## Supplementary Fig. 5 for "CHIMERA-DDR: A Machine Learning Framework for Classifying Heterogeneous Mismatch-Repair and Homologous-Recombination Deficiency Patterns in Prostate Cancer"

### Supplementary Figure 5

A

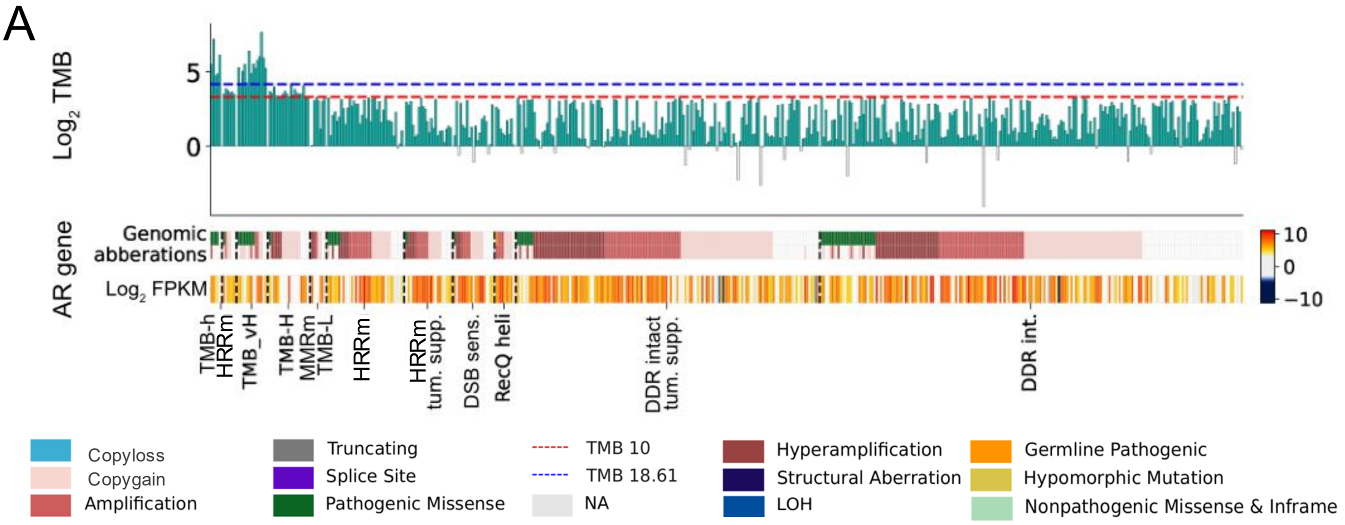


B

| AR gene mutation/ Amp | Mutation | No change | Odds_Ratio | p_value | Gain | Amp | Gain+Amp | Hemizygous | Odds_Ratio | p_value | Total Samples |
|---|---|---|---|---|---|---|---|---|---|---|---|
| TMB-vH HRRm | 4 | 1 | **26.923** | **0.001686827** | 0 | 1 | 1 | 4 | 0.106 | 0.033237941 | 5 |
| TMB-H HRRm | 1 | 5 | 1.346 | 0.572436602 | 3 | 1 | 4 | 2 | 0.851 | 1 | 6 |
| TMB-vH | 8 | 6 | **8.974** | **0.000268619** | 6 | 3 | 9 | 5 | 0.766 | 0.764117893 | 14 |
| TMB-H | 1 | 18 | 0.374 | 0.480673041 | 10 | 5 | 15 | 4 | 1.596 | 0.598270962 | 19 |
| MMRD-TMB-L | 0 | 7 | 0 | 0.599863823 | 2 | 3 | 5 | 2 | 1.064 | 1 | 7 |
| HRRm | 6 | 30 | 1.346 | 0.595870921 | 10 | 17 | 27 | 9 | 1.277 | 0.69112694 | 36 |
| HRRm tum. Supp. | 1 | 21 | 0.321 | 0.487256915 | 6 | 11 | 17 | 5 | 1.447 | 0.623908587 | 22 |
| DSB Sensor | 1 | 18 | 0.374 | 0.480673041 | 6 | 7 | 13 | 6 | 0.922 | 1 | 19 |
| RecQ Helicase | 1 | 8 | 0.841 | 1 | 4 | 4 | 8 | 1 | 0.162 | 0.45257431 | 9 |
| Intact tum supp | 8 | 136 | 0.396 | 0.027431345 | 48 | 72 | 120 | 24 | 2.128 | 0.005142879 | 144 |
| Intact | 26 | 175 | | | 66 | 75 | 141 | 60 | | | 201 |

C

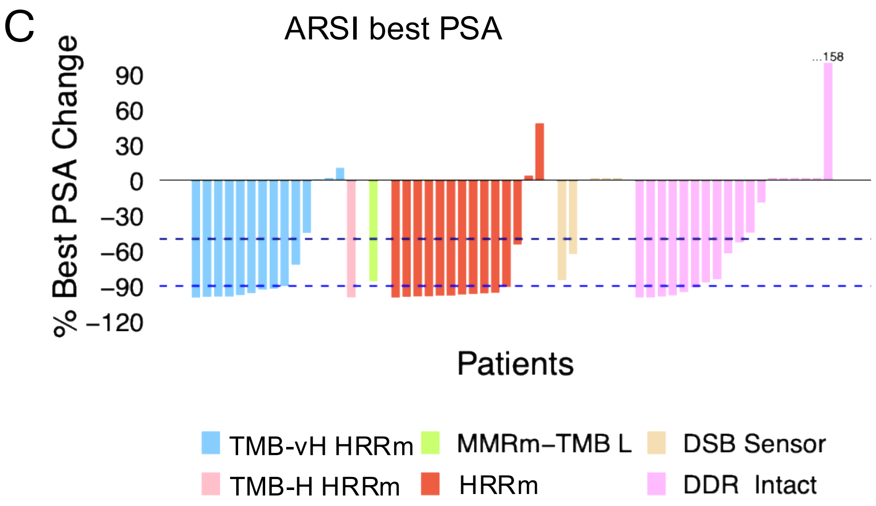
