## Supplementary Fig. 6 for "CHIMERA-DDR: A Machine Learning Framework for Classifying Heterogeneous Mismatch-Repair and Homologous-Recombination Deficiency Patterns in Prostate Cancer"

### Supplementary Figure 6

A

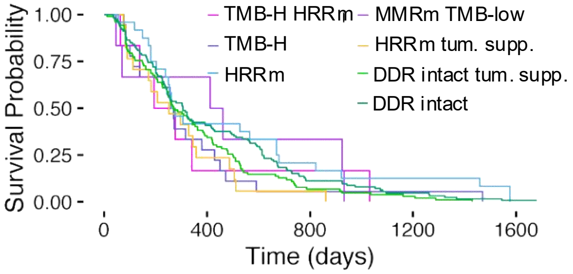


B

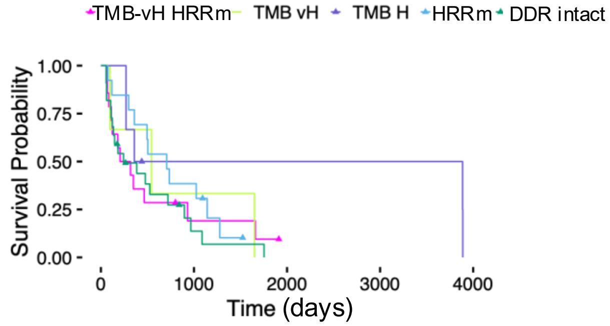


D

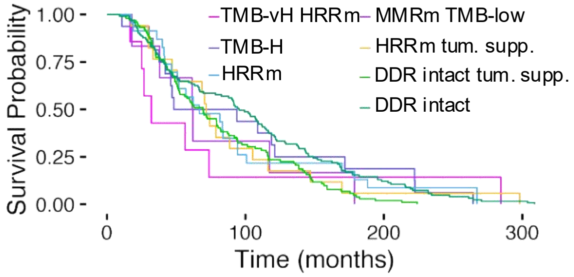


C

| | # samples | MSH2 | p_value | Odds_Ratio |
|---|---|---|---|---|
| TMB vH HRRm | 25 | 16 | **0.0000106** | 37.5 |
| TMB H HRRm | 8 | 1 | 0.444 | 3.14 |
| TMB vH | 19 | 9 | **0.00223** | 19.21 |
| TMB H | 29 | 11 | **0.00657** | 13.47 |
| HRRm | 17 | 0 | 1 | 0 |
| MMRm TMB Low | 1 | 0 | 1 | 0 |
| DDR intact | 24 | 1 | | |
| DSB sens. | 7 | 0 | 1 | 0 |

| | # samples | PTEN | p_value | Odds_Ratio |
|---|---|---|---|---|
| TMB vH HRRm | 25 | 3 | **0.0255** | 0.2 |
| TMB H HRRm | 8 | 0 | **0.0353** | 0 |
| TMB vH | 19 | 0 | **0.00107** | 0 |
| TMB H | 29 | 1 | **0.00123** | 0.05 |
| HRRm | 17 | 6 | 0.753 | 0.77 |
| MMRm TMB Low | 1 | 0 | 1 | 0 |
| DDR intact | 24 | 10 | | |
| DSB sens. | 7 | 3 | 1 | 1.05 |

| | # samples | SPOP | p_value | Odds_Ratio |
|---|---|---|---|---|
| TMB vH HRRm | 25 | 3 | 0.702 | 0.69 |
| TMB H HRRm | 8 | 0 | 0.55 | 0 |
| TMB vH | 19 | 0 | 0.118 | 0 |
| TMB H | 29 | 0 | **0.0363** | 0 |
| HRRm | 17 | 0 | 0.128 | 0 |
| MMRm TMB Low | 1 | 0 | 1 | 0 |
| DDR intact | 24 | 4 | | |
| DSB sens. | 7 | 0 | 0.55 | 0 |

E

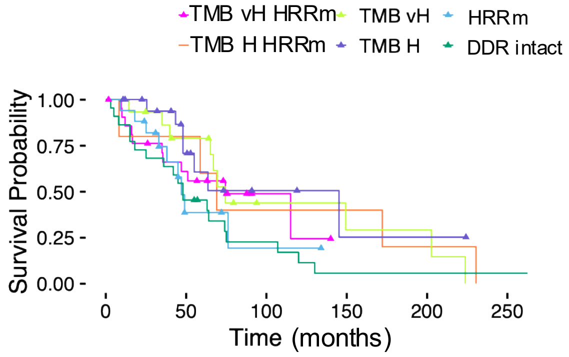


F

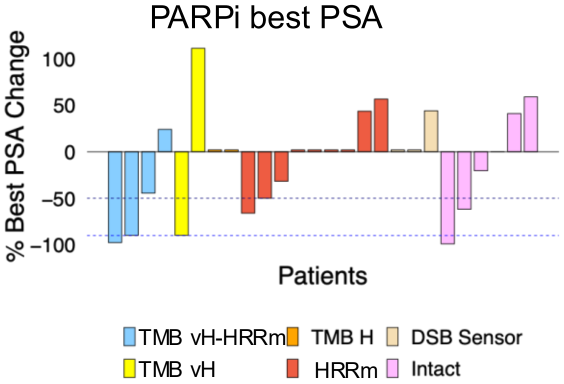
