## Supplementary Fig. 10 for "CHIMERA-DDR: A Machine Learning Framework for Classifying Heterogeneous Mismatch-Repair and Homologous-Recombination Deficiency Patterns in Prostate Cancer"

### Supplementary Figure 10

**Molecular subtyping based on traditional approach:**

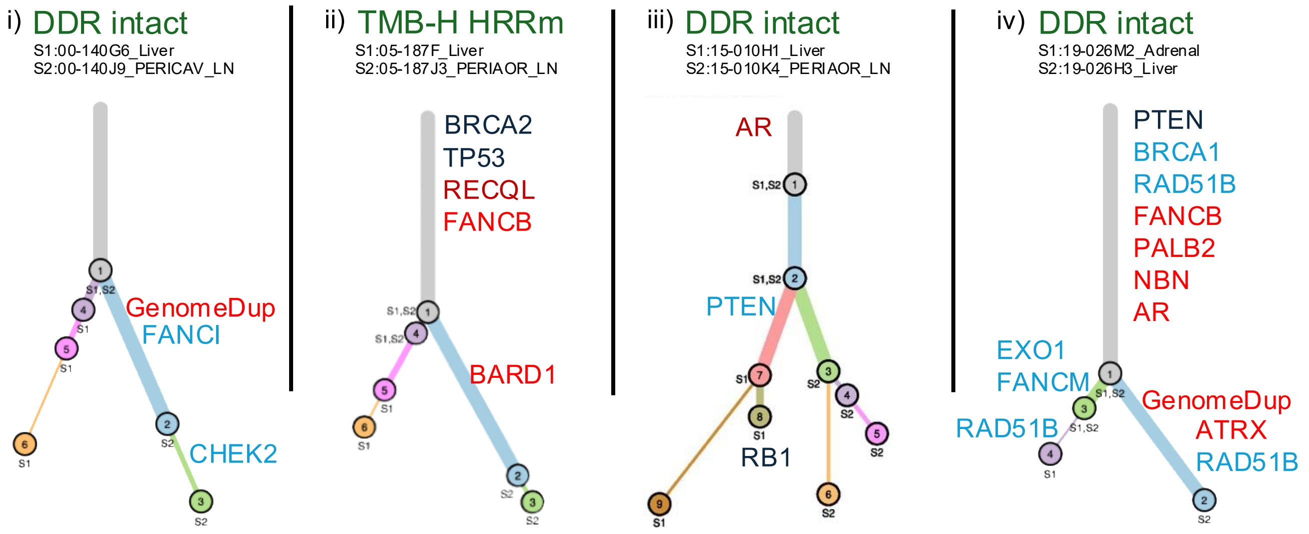


**CHIMERA-DDR predicted probabilistic subgroup annotation: post 2 tier analysis**

DDR intact

HRRd major

HRRd with clonal ambiguity (S1: Tum supp. mutated)

DDR intact tum supp.
