## Supplementary Figure Legends for "CHIMERA-DDR: A Machine Learning Framework for Classifying Heterogeneous Mismatch-Repair and Homologous-Recombination Deficiency Patterns in Prostate Cancer"

**Supplementary Figure 1:** Schematics showing research methodology workflow illustrating the progression from recognizing limitations in binary classification for patient stratification through the creation of CHIMERA-DDR tool to adopted validation approaches.

**Supplementary Figure 2:** A) Allele-specific percent genome alteration analysis in the 482 exome cohort demonstrates distinct PGA patterns in allele A and allele B within the TMB-High subgroup, revealing differential genomic instability between TMB-vH HRRm and TMB-H HRRm tumor subsets. B) Scatter plot demonstrating the relationship between MSI-high scores and MMRd signature scores. Analysis reveals that MSI-H tumors are almost invariably MMRd signature positive, while MMRd signature-positive tumors may not necessarily exhibit MSI-high status. C) Scatter plot demonstrating the relationship between MSI-high scores and tumor mutational burden (TMB). Analysis reveals that tumors with >18.62 mutations/Mb are significantly enriched for MSI-high status. D) Scatter plot demonstrating the relationship between TMB and COSMIC MMRd signature scores. Analysis reveals that tumors with >18.61 mutations/Mb are almost always MMRd signature positive. E) Three-fold cross-validation analysis demonstrates optimal TMB cutoff determination through AUC-ROC curves from binary logistic regression. Results indicate that 18.61 mutations/Mb represents the optimal threshold above which maximum enrichment for MSI-high status is observed. F) Gene expression analysis demonstrates significant MSH2 downregulation in TMB-vH and TMB-H tumor subsets. G) TP53 pathogenic mutation analysis reveals borderline significant enrichment of TP53 mutations in the DSB sensor subset (comprising tumors with ATM, CHEK2, DNA2, EXO2 mutations) compared to the DDR-intact subset. No statistical enrichment was observed in other DDR subgroups. This analysis was performed with tumor groupings agnostic to TP53 mutation status (tumor suppressor mutation status was excluded from subset definitions).

**Supplementary Figure 3:** A) Subgroup pooled and subsampled (300X) copy number analysis using CNVkit demonstrates recurrent copy number patterns across 9 of the 11 DDR axis subsets. RecQ helicase and DSB sensor gene mutant subgroup profiles are not shown. Analysis revealed absence of AR gene and 8q gain/amplification in TMB-vH and TMB-vH HRRm subsets, which are otherwise highly recurrent events in other tumor subgroups. B) Recurrent copy number instability enrichment across 11 DDR axis subgroups.

**Supplementary Figure 4:** A) Component bar plot displaying average immune infiltrate cell frequencies using CIBERSORT absolute mode analysis. Subgroup means represent average immune cell infiltrate signatures per group, revealing relatively high CD8 T cell frequencies in TMB-vH HRRm and TMB-vH tumor subsets. Notably, the TMB-H group is also showing a substantially elevated frequency of CD8 T cells. B) Comparative gene expression profiles of immune checkpoint pathway genes across all 11 DDR axis subgroups. This analysis expands upon Fig. 3G, which displayed only 5 DDR mutant groups, to include comprehensive representation of all defined subgroups. C) Heatmap displaying expression of 46 selected DDR and DDR-associated tumor suppressor and oncogenes across 462 advanced prostate cancer tumors, including tumor prognostic scores (Top panel, CCP score, NE activity score, and AR activity scores). Notably, MMR pathway gene overexpression (red) is absent in TMB-vH and TMB-vH HRRm subsets, which is otherwise observed in at least limited samples across all other tumor subgroups. Instead, a subset of TMB-vH and TMB-vH HRRm tumors exhibit MSH2 low expression (blue).

**Supplementary Figure 5:** A) Heatmap showing the relationship between TMB and AR gene expression alongside AR gene genomic status (pathogenic mutations and copy number aberrations). The analysis demonstrates high frequency of AR pathogenic mutations and limited copy number gain/amplification in TMB-vH and TMB-vH HRRm tumor subsets. A positive correlation between AR gene expression and AR gene amplification/gain is evident. Interestingly, other DDR subgroups show no association between AR amplification or mutations and TMB patterns below 10 mutations/Mb; instead, TMB distributions appear random. The HRRm subset consistently exhibits higher TMB values approaching 10 mutations/Mb, while other subsets display more variable TMB ranges. B) Table confirming the observations from Supplementary Fig. 4A, demonstrating positive enrichment of AR pathogenic mutations in TMB-vH and TMB-vH HRRm subsets compared to the DDR-intact advanced prostate cancer subset, while showing negative enrichment of AR gene copy number gain/amplification. C) Swimmer plot displaying best PSA response compared to baseline in advanced prostate cancer patients stratified by DDR axis subgroups receiving ARPI treatment. Analysis reveals a trend toward more frequent PSA90 response attainment in TMB-vH HRRm and HRRm tumors compared to DDR-intact tumors in the clinical validation cohort of 130 cases. Notably, ARPI treatment data was unavailable for the majority of other patients, resulting in underrepresentation of other DDR subgroups.

**Supplementary Figure 6:** Clinical cohort analysis revalidated discovery cohort findings, confirming distinct molecular patterns between TMB-vH and TMB-H subsets. A & B) Kaplan-Meier plots showing ARSI response where TMB-vH tumors did not demonstrate statistically significant differences in treatment duration patterns or trends. A) ARSI response in SU2C discovery cohort grouped according to 11 DDR subgroup definitions. B) Validation Kaplan-Meier analysis for time on treatment with ARSI. For clarity, we are showing 7-group and 4-group results in A and B, respectively. C) Mutation frequency analysis demonstrating significant enrichment of MSH2, PTEN, and SPOP alterations in the clinical validation cohort, corroborating discovery cohort findings and further establishing the distinct molecular profile of the TMB-vH subgroup. D & E) Kaplan-Meier analysis of overall survival from diagnosis demonstrates TMB-vH as a prognostically distinct subgroup. D) Overall survival analysis in the discovery cohort (482 patients) showing TMB-vH with HRR mutations trending toward worse outcomes (magenta). E) Validation cohort analysis where TMB-vH with HRR mutations (magenta) trended toward relatively better outcomes compared to other subgroups. Notably, discovery cohort patients rarely received ICI or PARPi therapy, whereas the validation cohort had broader access to these emerging mainstream therapies, potentially explaining the improved outcomes observed in the TMB-vH HRRm subgroup. F) Swimmer plot showing nonspecific PSA50 outcome in validation cohort patients stratified based on 11 DDR subgroup definitions.

**Supplementary Figure 7:** UW-TAN exome cohort (n=270 tumors from 112 individual patients) intratumoral heterogeneity determination across 4 DDR genomic signatures. A) Showing relatively rare intratumoral heterogeneity in terms of MSI signature scores. TMB demonstrated relatively greater intratumoral heterogeneity, although we noted limited instances where intratumoral scores crossed TMB-H or TMB-vH cutoff limits, suggesting sample-specific ambiguity in patient identity determination. Analysis also revealed relatively greater intratumoral heterogeneity in terms of HRD-Scar scores and LOH scores. B) Subtype-specific analysis of metastatic sample tissue origin showed relatively even distribution across multiple subtypes. Metastatic tissue sampling sites demonstrated limited delineation in TMB-vH HRRm and TMB-H HRRm, MMRm TMB-Low, and other rare subgroups. We inferred that lower representative sample numbers in these respective classes are the potential reason for such observed bias toward specific tissue sampling sites.

**Supplementary Figure 8:** CHIMERA performance determinants and validation. A) Distribution of tumor cellularity in 445 tumors used in CHIMERA training. Notably, tumors with TMB-vH HRRm or TMB-H HRRm, those with mutations in DSB sensor pathways, or tumors with MMR gene bi-allelic mutations but <10 TMB were kept on hold (n=37 tumors) for independent testing post-CHIMERA training and cross-validation. B) DDR-intact-like predicted probability in tumors with 7 CHIMERA-DDR class assignments following 2-tier analysis. Analysis suggests DDR-intact predicted probability is not a function of sample tumor cellularity but rather is primarily guided by the tumor's DDR-intact-like molecular characteristics.

**Supplementary Figure 9:** CHIMERA outcome showing distribution of predicted probabilities across 7 prediction class components in tumors with 5 distinctive DDR classes. Note: Class identity assignment prior to CHIMERA-DDR training were done based on their curated detection of genomic aberrations.

**Supplementary Figure 10:** PhyloWGS-based clonal evolution analysis of DDR pathway alterations in tumor samples. PhyloWGS, a computational method for tumor evolution analysis that reconstructs genetic lineage of cancer cells from whole-genome sequencing (WGS) data, was used to reconstruct clone-aware DDR genomic aberration events in 4 patients. All patients had paired WGS data from two tumor samples. (Green font- pre-analysis genomics-based assigned molecular subtype.) Blue font- CHIMERA-DDR predicted assigned class. Branch length: Represents the relative mutational burden or evolutionary distance (Relative frequency of somatic events). Node width: Indicates clonal prevalence (cellular fraction that carries those somatic events).

i) Patient 00-140 – DDR-intact:CHIMERA-DDR-intact: DDR-intact characteristics are supported by clone-aware assessment of mutations and copy numbers. Pericaval tumor harbors FANCI heterozygous loss and subclonal loss of monoallelic CHEK2. Overall, clonal architecture of patient 00-140 supports DDR-intact inference.

ii) Patient 05-187 - TMB-H HRRm::CHIMERA-HRRd: Tumor exhibits TMB ~14 mut/Mb, lacks MSI signature scores, and shows absent COSMIC MMRD signature scores. BRCA2 bi-allelic loss-of-function is clonal, and the detected mutation (p.N2135Kfs*32) is pathogenic with high penetrance for HRRd phenotype.

iii) DDR-intact::CHIMERA-HRRd with ambiguous dominant molecular type inference: Demonstrates relatively elevated subclonal evolution compared to other studied tumors. One of two tumors showed >0.2 HRRd probability while the other showed 0.13 probability of HRRd with tumor suppressor mutations. Clonal analysis detected subclonal heterozygous loss of PTEN and rare clonal loss of RB1. No homologous recombination pathway gene aberrations were detected that passed quality control thresholds.

iv) DDR-intact::CHIMERA-HRRd: Clone-aware DDR genomic analysis revealed clonal heterozygous loss of BRCA1 and RAD51B, with subsequent independent subclonal copy-neutral loss of heterozygosity events in RAD51B. Despite input annotation as DDR-intact, CHIMERA inferred HRRd-major classification, suggesting functional homologous recombination loss-of-function.
